# Midday shading alleviates thermal stress but has contrasting agronomic effects across medicinal and aromatic plants

**DOI:** 10.64898/2026.09.18.752694

**Authors:** Ombeline Decuy, Benjamin Lemaire, Cyril Bozonnet, Soline Morel, Clara Paredes, Laurent Barroux, Pascal Walser, Philippe Balandier, Stephane Herbette

## Abstract

Climate change is increasing the frequency and intensity of heatwaves and droughts, threatening medicinal and aromatic plants (MAP) production. Temporary shading during the hottest part of the day may reduce heat and radiative stress while limiting the reduction in light availability. We evaluated this strategy in a field experiment using three MAP species with contrasting ecological requirements and harvested organs: lavandin (*Lavandula × intermedia*), curly parsley (*Petroselinum crispum*) and valerian (*Valeriana officinalis*). Vertical shade nets provided midday shading and effects on crop microclimate, leaf temperature, water status, PSII photochemical efficiency, plant growth and biomass and essential oil production were assessed over two growing seasons. Midday shading reduced daily photosynthetically active radiation by 19.9-55.9%, with weak effects on air temperature and vapor pressure deficit. In contrast, leaf temperature at solar noon decreased by 6.9 °C, 5.6 °C and 3.0 °C in lavandin, valerian and parsley, respectively. Midday shading improved light-adapted and maximum PSII photochemical efficiencies in all three species and reduced branch dehydration in lavandin during heatwave periods. However, agronomic responses were species-dependent. Floral biomass decreased in lavandin, whereas parsley leaf biomass and valerian root biomass increased. Essential oil concentration was unaffected in lavandin and valerian. Overall, our results show that midday shading primarily reduced leaf-level thermal and radiative stress rather than modifying the surrounding air microclimate, and suggest that its agronomic effects may reflect a balance between stress alleviation and reduced carbon acquisition, that varies with species ecology and production objectives.

## 1. Introduction

Medicinal, aromatic and perfume plants (MAPs) comprise a highly diverse group of several thousand cultivated or wild-harvested species, including more than 2,000 species found in Europe (Lubbe and Verpoorte, 2011; Pandey et al., 2020). They play a major economic, social, cultural and ecological role worldwide, supporting local communities and supplying raw materials for numerous industrial sectors (Pergola et al., 2024). They include annual, biennial and perennial herbs, shrubs and trees originating from a wide range of climatic regions. MAPs are valued for diverse harvested organs, including flowers, leaves, roots, seeds and bark, which are processed into essential oils, pharmaceuticals, perfume, cosmetics, culinary ingredients and many other products (Lubbe and Verpoorte, 2011; Rao et al., 2024). Unlike conventional crops, the agronomic value of MAPs depends not only on biomass production but also on the yield and chemical composition of their secondary metabolites, which determine the quality and commercial value of the final products (Figueiredo et al., 2008; Lubbe and Verpoorte, 2011). Like other agricultural sectors, MAP cultivation is increasingly threatened by climate change. In Europe, rising temperatures, more frequent and intense heatwaves and changes in precipitation patterns are posing major challenges for crop production systems (Calvin et al., 2023). Abiotic pressures, like heat and drought, are the leading cause of crop failure worldwide, reducing the average yields of major crops by more than 50% (Bray et al., 2000). While no projections of future climate change impacts on MAP yields are currently available, projections for major cereal crops highlight the potential threat to agricultural productivity, with yield losses ranging from 6.2% to 18.3% in the absence of adaptation strategies.(Rezaei et al., 2023).

In this context, cultivating crops under shaded conditions may be a promising strategy to mitigate the impacts of climate change (El-Zawily et al., 2024; Gosme et al., 2016).Shading modifies the crop microclimate by reducing incident solar radiation, air and plant surface temperature. Depending on environmental conditions, Shading can also increase air relative humidity and reduce evaporative demand (Jose, 2009; Mahmood et al., 2018; Ukwu et al., 2025). Various shading strategies have been developed, including shade nets, agrivoltaic systems and agroforestry (Juillion et al., 2022; Mahmood et al., 2018; Monteith et al., 1991). Conventional shade nets generally provide a homogeneous and continuous reduction in irradiance, with different shading levels and spectral properties depending on the net characteristics. They generally reduce air, leaf and fruit temperatures, although their effectiveness depends on ventilation, and poorly ventilated systems may occasionally promote heat accumulation (Mupambi et al., 2018; Narjesi et al., 2023; Tezcan et al., 2022). In contrast, agrivoltaic and agroforestry systems create heterogeneous and dynamic light environments in which the magnitude and duration of shading vary according to canopy architecture and, crop position relative to the solar trajectory (Dupraz et al., 2018; Mantino et al., 2021; Pallotti et al., 2023; Touil et al., 2021; Zhao et al., 2003). Consequently, the resulting microclimatic conditions differ among systems, with the strongest cooling effects generally occurring during periods of high atmospheric demand (Ali Abaker Omer et al., 2025; Juillion et al., 2022; Pang et al., 2019; Ukwu et al., 2025). More recently, vertical shade nets have been proposed to reproduce the partial shading conditions made by trees in agroforestry systems. Although their effects on plant growth and physiology have begun to be investigated, their impact on crop microclimate remains largely unexplored (Zubay et al., 2021).

Shading affects plant performance through two contrasting effects. On one hand, the reduction in photosynthetically active radiation (PAR) under shaded conditions induces a range of morphological and physiological adjustments in plants. A common response is a decrease in leaf mass per area (LMA: (Poorter et al., 2009) to maximize light interception per unit of biomass invested, and thus enhanced photosynthetic efficiency when light becomes a limiting resource (Arenas-Corraliza et al., 2019; Poorter et al., 2009). However, prolonged or excessive shading may reduce photosynthesis, growth, biomass production and yields due to carbon limitation (Inurreta-Aguirre et al., 2018; Querné et al., 2017; Temani et al., 2021). Shading can cause a yield loss of approximately 10 to 30%, depending on crops, with losses exceeding 50% under dense, unmanaged tree canopies (Muschler, 2016; Temani et al., 2021; Weselek et al., 2021). On the other hand, moderate shading can alleviate thermal, hydric and radiative stress by reducing leaf overheating, photoinhibition, crop evapotranspiration and plant water deficit. By lowering excess irradiance reaching the photosynthetic apparatus, moderate shading has been shown to maintain higher PSII efficiency than full sunlight, as reflected by higher values of both the PSII efficiency (F’_v_/F’_m_ under ambient light and the maximum quantum efficiency of PSII (F_v_/F_m_) after dark adaptation, indicating a lower degree of photoinhibition (Alam et al., 2018; Baker, 2008; Chichaghare et al., 2026; Magarelli et al., 2025; Zha et al., 2022). In parallel, the reduction in evaporative demand may improve plant water status, which is commonly assessed through water potential (Ψ), an integrative indicator of plant water deficit (Kramer, 1988; Ziegler et al., 2024). Evidence from shade-net, agrivoltaic and agroforestry systems shows that shaded plants maintain less negative midday water potentials (Ψ*_md_*) than plants grown under full sunlight, reflecting an improved plant water status (Bernal-Basurco, 2026; Kabir et al., 2022; Magarelli et al., 2025; Mira-García et al., 2022; Montanaro et al., 2009). Under hot and dry conditions, these physiological benefits may compensate for the reduction in light availability, and several studies conducted in agroforestry and agrivoltaic systems have reported maintained or even increased crop yields compared with full sunlight, highlighting the importance of stress alleviation under conditions of high atmospheric demand (Barron-Gafford et al., 2019; Blanchet et al., 2022; Jha et al., 2026; Weselek et al., 2021) The effect of shading would therefore depend on the balance between reduced carbon acquisition under low irradiance and the alleviation of heat and water stress.

Around solar noon, when net radiation and atmospheric evaporative demand reach their daily maxima, plants experience their greatest surface temperature and lowest leaf water potential (Ψ), often resulting in stomatal closure and midday depression of photosynthesis (Betts, 2009; Hirasawa and Hsiao, 1999; Xiao et al., 2021). We thus assume that temporary shading restricted to the most stressful hours of the day may represent an effective compromise, providing protection against heat and water stress while maintaining sufficient irradiance to sustain plant growth and yields.

The effects of shading on MAPs are species-dependent. While several MAP species maintain satisfactory yields (biomass and essential oil) and product quality under shaded conditions, some species may even benefit from moderate shading (30%) (Lalević et al., 2023; Zubay et al., 2021). In contrast, reduced light availability has been associated with significant yield losses in more heliophilous species (Lalević et al., 2023; Palada et al., 2004; Rao et al., 2004; Singh et al., 1998). For example, Mambrí et al., (2018) found that a 50% shade net reduced flower yield by 37% and essential oil yield by 32 to 46% in *Lavandula dentata*. These contrasting responses likely reflect differences in species ecology and adaptation to contrasting light environments, as the morphological and physiological traits that enhance performance under low irradiance are often incompatible with those required to cope with high irradiance, heat and desiccation (Valladares and Niinemets, 2008). In addition, agronomic consequences of shading are likely to depend on the harvested organ, as light availability influence carbon partitioning. Under low-light conditions, plants generally allocate a greater proportion of biomass to leaves to maximize light interception often at the expense of root growth (Poorter et al., 2012). At the same time, reduced irradiance tends to decrease flower and seed production by limiting biomass accumulation and shifting resource allocation towards vegetative organs (Marcelis, 1996; Van Baalen et al., 1990). These findings are almost exclusively based on permanent shading conditions, and the physiological and agronomic responses of MAPs to temporary midday shading remain largely unexplored (Katsoulis et al., 2022; Lalević et al., 2023; Zubay et al., 2021).

The objective of this study was (i) to quantify the microclimatic changes induced by midday shading and (ii) to determine how these changes affect the MAPs physiology and yields in three contrasting MAP species. To specifically investigate the effects of temporary midday shading, we developed an artificial shading system consisting of vertical shade nets positioned on the southern side of the crop rows, providing shade around solar noon.

To asses the potential of midday shading as an adaptation strategy for the MAP sector, three contrasting MAP species, representative of the sector and differing in ecology, harvested organs and life cycle were selected: lavandin (*Lavandula x intermedia* Emeric ex Loisel), curly parsley (*Petroselinum crispum* Mill.) and valerian (*Valeriana officinalis* L.).

We hypothesized that (i) partial shading applied during the hottest hours of the day would attenuate peak incident radiation, air temperature and vapour pressure deficit, thereby buffering the crop microclimate during periods of maximum atmospheric constraint; (ii) midday shading would thus reduce plant water stress and improve physiological functioning; (iii) the alleviation of thermal and water stress would outweigh the reduction in irradiance, resulting in contrasting agronomic responses depending on species and harvested organ; and (iv) flower production would be negatively affected in the heliophilous lavandin, whereas leaf biomass in parsley and root biomass in valerian, which is a shade-tolerant species, would be maintained or enhanced under midday shading.

To test these hypotheses, we compared the effects of midday shading on the crop microclimate, plant physiological responses and agronomic performance. Microclimatic measurements included incident PAR, sheltered air temperature, and VPD. The impact of the modified radiative environment on the plants was assessed through measurements of leaf surface temperature. Plant responses were evaluated by measuring leaf mass per area (LMA), shoot Ψ*, and maximal and light-adpated photochemical efficiencies* (F_v_/F_m_ and F′_v_/F′_m_). Continuous branch diameter monitoring using LVDT in lavandin was used to quantify radial growth and maximum daily shrinkage (MDS) as an integrative indicator of plant water status (Fernández and Cuevas, 2010; Lamacque et al., 2020; Ortuño et al., 2010). Agronomic performance was assessed through measurements of biomass and essential oil yields.

## 2. Material and methods

### 2.1. Experimental site

The experiment was conducted in southern France (44.546236° N, 4.831945° E), an area under Mediterranean climate with strong seasonal contrasts with hot, dry summers and mild, wet winters. In this region, summer is marked by scarce precipitation and drought periods extending for several months. Most rainfall occurs in autumn and winter and may be intense and episodic.

The soil was a sandy loam, with 18.5 ± 1.1*%* clay, 22.5 ± 1.2*%* silt, and 58.9 ± 2.1*%* sand, with an alkaline pH of 8.37 ± 0.04. The soil had a medium cation exchange capacity (CEC) of 13.9 ± 1 *cmol kg ¹* and an effective depth of approximately 70 *cm*. The carbon-to-nitrogen (C/N) ratio was relatively low (5.4 ± 3.2), indicating a rapid organic matter turnover. All soil analysis data are provided in the supplemental data.

The experimental plot layout is shown schematically in Figure 1. Shading was achieved using vertical green/black shading nets providing 78% to 85% shade (FILET ALPHAOMBRE 70-7071-600KLY, Alphatex, France). The incident red-to-far-red (R/FR) ratio was 1.22 and decreased only slightly to 1.01 beneath the shade nets, indicating that the spectral composition of the incident radiation was only moderately affected by the shading treatment. Simulations were previously performed using the SAMSARALight model on the simulation capsis platform (Ligot *et al*., 2014) to determine the optimal dimensions and placement of the shading nets. The temporary shading was then achieved using nets of 2 *m* in height and positioned 70 cm from the central part of the plant and 30 *cm* above ground level to simulate a tree hedge configuration, increase the final height of the nets and allow the wind flowing below the net. The nets were oriented along an east–west axis, thereby providing temporary shading to the plants, around solar noon, during the hottest hours of the day. The shading structures were supported by 3 *m* high pine posts (8 cm in diameter), embedded 70 cm into the soil and spaced at 2 *m* intervals. The nets were tensioned and secured to the posts using 2 *mm* diameter galvanized steel cables running along the upper and lower edges. To enhance structural stability under wind load, two additional UV-stabilized high-density polyethylene (HDPE) cables (4 *mm* in diameter) were fixed directly to the posts on each side of the net at heights corresponding to one-third (0.66 *m*) and two-thirds (1.33 *m*) of the net height. The shading system was installed on 14 June 2024 and it was removed during the winter period, from 2 December 2024 to 8 April 2025, to prevent damage from winter windstorms and to mimic the absence of foliage in deciduous trees during winter.

**Fig. 1.**
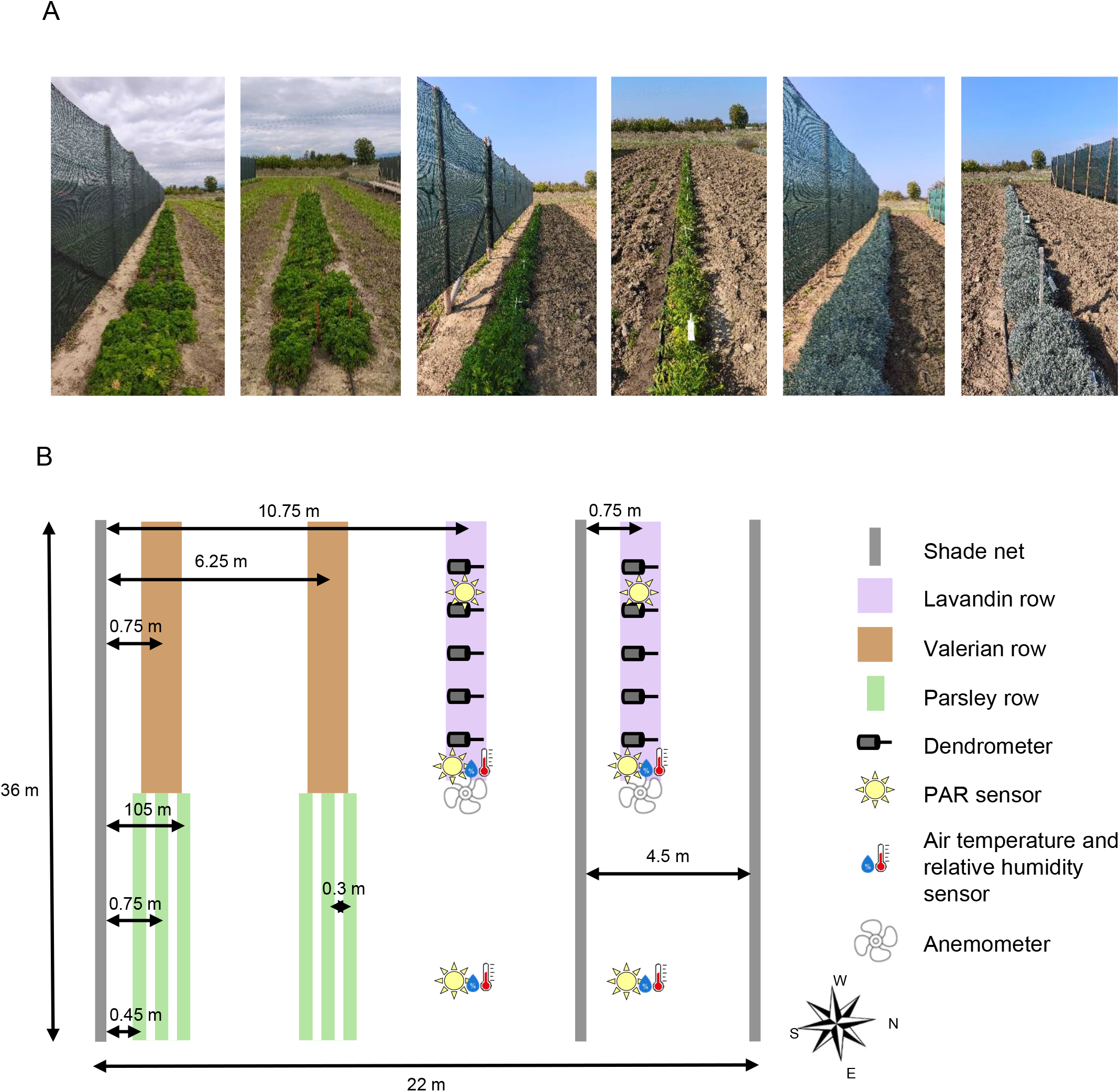
Photographs (A) and schematic representation (B) of the experimental plot. A) Photographs of the experimental setup illustrating the different plant species under shaded and control treatments. From left to right: shaded and control parsley plants, shaded and control valerian plants, and shaded and control lavandin plants. B) Schematic representation of the experimental plot indicating the distances between the shading nets and the center of the crop rows. Anemometer and PAR, air temperature, and relative humidity sensors were positioned at the center of the crop rows. Further details regarding sensor characteristics and distances are provided in the Materials and Methods section.

Three contrasting medicinal, aromatic and perfume (MAPs) species were selected as representative species of the sector: lavandin (*Lavandula × intermedia* Emeric ex Loisel), curly parsley (*Petroselinum crispum* Mill.) and valerian (*Valeriana officinalis* L.). These species were chosen to encompass a broad range of ecological preferences, life cycles and harvested organs representative of the diversity of MAP crops. Lavandin is a perennial Mediterranean species naturally adapted to warm, high-radiation environments, whereas parsley and valerian are temperate species with higher water requirements and contrasting tolerances to heat and shade. In addition, the three species differ in the harvested organ, with flowers collected for essential oil production in lavandin, leaves harvested in parsley and roots harvested in valerian. This diversity allowed the effects of midday shading to be evaluated across contrasting ecological strategies and production objectives.

Lavandin (*Lavandula × intermedia* Emeric ex Loisel) is one of the most important MAP crops in France. Lavandin flowers are harvested to extract essential oil. It is a perennial mediterranean perfume plant well adapted to environments characterized by high solar radiation and limited water availability (Lis-Balchin, 2002; Pokajewicz et al., 2023). The experiment was conducted within its optimal cultivation area. The cultivar Grosso ADA was established in a single row on 3 March 2024 using nine-month-old bare-root plants, with an inter-plant spacing of 45 *cm*. The lavandin row was 19 *m* long. Twenty days after planting, a mineral fertilization was manually applied at a rate of 50 *kg ha ¹* nitrogen (N), 50 *kg ha ¹* phosphorus (P), and 100 *kg ha ¹* potassium (K). In 2024, regular irrigation was not required due to regular rainfall and deep soil. In 2025, lavandin plants were irrigated on five occasions during July and August using a drip irrigation system, with an average irrigation duration of 2 *h* 50 *min* per event and a flow rate of 1.6 *L .h ¹* from drippers spaced 33 cm apart.

Valerian (*Valeriana officinalis* L.) is a medicinal plant native to the temperate regions of Europe and Asia. The roots are used for their sedative and anxiolytic effects. It typically grows in moist habitats, like woodlands and meadows, under full sun to partial shade and is adapted to cool, temperate climates with consistently high soil moisture (Lee et al., 1996; Patočka and Jakl, 2010). Accordingly, the experiment was conducted under warmer and drier climatic conditions than those typically encountered across its cultivation range. The cultivar VALIA was established from plug seedlings, sown annually in February and transplanted on 3 June 2024 and 19 May 2025, with an inter-plant spacing of 30 *cm*. Valerian row length was 13 *m* for plants transplanted in 2024 and 18 *m* for those transplanted in 2025. In 2025, a mineral fertilization supplying 50 *kg ha* ¹ N, 70 *kg ha ¹* P, and 150 *kg ha ¹* K was applied manually to the soil on 11 June, approximately one month after transplanting. In 2024, valerian was irrigated using drip irrigation on seven occasions, including six events during July and August and one in September, with an average irrigation of 44 *mm.* In 2025, irrigation was applied one to three times per week from May to August, depending on temperature and rainfall conditions, for a total of 21 irrigation events, with an average irrigation of 34 *mm* per event.

Curly parsley (*Petroselinum crispum* (Mill.) var. *crispum*) is an aromatic species cultivated worldwide in temperate regions for its edible leaves. It grows optimally under moderate climatic conditions (Brengi et Nasef, 2023; Marthe, 2020). It is generally cultivated in full sun, although it can also grow in half-shady environment, prefers moist soils, and it is relatively tolerant to low temperatures, whereas high temperatures may reduce growth (Agyare et al., 2017; Brengi and N. Nasef, 2023; Marthe, 2020). The experiment was conducted under warmer and drier climatic conditions than those prevailing within its usual cultivation range. This aromatic herb is a biennial plant that is grown annually. The crop can be harvested several times during the year before being destroyed (Marthe, 2020;). The above-ground part of the plant is harvested for sale fresh, dried, or frozen (Sabry et al., 2016). The cultivar Bravour was sown in three rows at a density of 125 seeds per linear meter on 26 June 2024 and 10 April 2025. Parsley rows were 13 *m* long and were established at 0.45, 0.75, and 1.05 *m* from the net. After each harvest, except the final cut, parsley plants were fertilized manually with a mineral fertilizer supplying 50 *kg ha ¹* N, *80 kg ha ¹* P, and 200 *kg ha ¹* K. In 2024, parsley was irrigated using drip irrigation on seven occasions between June and September, depending on rainfall and temperature conditions, with an average irrigation of 48 *mm.* In 2025, irrigation was applied one to three times per week from May to August according to weather conditions, for a total of 26 irrigation events from sowing to September, with an average irrigation duration of 2 *h* 30 *min* per event and a flow rate of 1.6 *L .h ¹*.

Control plots were established using identical planting and sowing procedures but without shading nets. Control rows of valerian, parsley, and lavandin were located at a sufficient distance from any net (≥ 6.25 m) to avoid any shading, as determined by simulations using the SAMSARALight model and checked throughout the seasons (Fig.1). For all three crops, fertilization and irrigation regimes were identical between midday-shaded and control conditions; midday-shading was the only factor differing between treatments.

### 2.2. Microclimate parameters

To assess the effect of shading nets on the wind regime, wind speed and direction were measured using combined anemometers and vanes (Davis Instruments, USA), installed both behind the shading net and in the control (unshaded) condition (Fig. 1). The sensors were positioned above the lavandin canopy.

Photosynthetically Active Radiation (PAR, 400 – 700 *nm*) was measured using three PAR/CBE 80 sensors (SOLEMS, FR) per modality. PAR sensors were installed on the lavandin row located 0.75 *m* from the net, at 50 *cm* above ground level (Fig. 1), i.e. at canopy height.

Air temperature and relative humidity were monitored using external hygrometry (±1.5*%*) and temperature (±0.1 *°C*) sensors housed in METSPEC RAD14 weather shelter (Campbell Scientific, FR). Sensors were positioned on the lavandin row at canopy height.

The vapor pressure deficit (VPD) was calculated from air temperature and relative humidity data using the following equation:

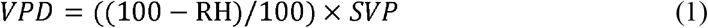

where VPD is the vapor pressure deficit (*Pa*), RH is relative humidity (*%*) and SVP (*Pa*) is saturation vapor pressure. For convenience VPD is given in kilopascals (*kPa*). The saturation vapor pressure was calculated according to the Tetens equation:

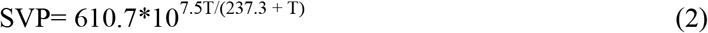

Where T is temperature (*°C*) (Monteith and Unsworth, 2008; Murray, 1967).

The spatial arrangement and positioning of all sensors are shown in Figure 1. All sensors were connected to centralized data acquisition units (e-PépiPIAF 1.1 dataloggers, Capt Connect, Clermont-Ferrand, France) housed in waterproof IP67-rated enclosures. Measurements were recorded every minute, and hourly means were subsequently calculated for data analysis.

### 2.3. Surface temperature

Thermal images have been acquired on 16 July 2025 at solar noon and on 17 July 2025 in the morning before shade (full sun days) using a FLIR thermal camera (model T650sc) positioned above the apex of the plant of interest. Just prior to image acquisition, a black sheet was positioned on the soil around the plant to obtain a homogeneous background. Resulting images were then treated in two steps. First, using an existing Python script (https://github.com/ITVRoC/FlirImageExtractor) that extracts raw radiative data from the image metadata and convert them to pixel-scale temperature values. In a second step, these temperatures maps were analyzed using a custom ImageJ macro (Schindelin et al., 2012) where the user is first asked to draw a box around the plant of interest that is used as prompt to SAMJ [https://arxiv.org/abs/2506.02783] in order to extract the plant contours. The user is additionally asked to provide an optional threshold to exclude extreme temperatures values that belong to the soil. The resulting mask is used to extract the mean plant temperature and the temperature standard deviation. The macro can work on a batch of image. These tools are freely available on GitHub: https://github.com/cyrilbz/thermal_images.

### 2.4. Leaf mass area

Leaf mass per area (LMA) measurements were performed on lavandin plants from 23 to 26 August 2024 and from 29 September to 1 October 2025. For each plant and treatment, six fully expanded current-year leaves were collected from nodes 4, 5, and 6 (counted from the apex), with a total of twelve plants per treatment in 2024 and nine in 2025. For valerian, LMA measurements were conducted from 23 to 25 September 2025. One fully expanded mature leaf located at the outer part of the rosette was collected per plant per modality, with a total of twelve plants per treatment.

For both species, after harvest fresh leaves were immediately placed in a sealed plastic bag and brought to the laboratory. Then, they were scanned using a flatbed scanner, and leaf area was determined using ImageJ software (National Institutes of Health, Bethesda, MD, USA). After scanning, leaves were oven-dried at 100*°C* for at least 3 h until constant weight to determine leaf dry mass with a precision balance. Leaf mass area (LMA) was calculated as the ratio of leaf dry mass to leaf area (*g.m ²*) (Poorter et al., 2009).

LA and LMA could not be determined for curly parsley because its highly dissected and curled leaves made accurate leaf area measurements technically challenging.

### 2.5. Stem diameter variations

Stem diameter variations were only measured in Lavandin as the use of dendrometers required a woody stem. From the 1^st^ of July 2024 to the end of the experiment, branch diameter was continuously measured using miniature displacement sensors with a friction free core glued to the bark and LVDT (model DF2.5, DF5.0 and MD5; Solartron Metrology, Massy, France) connected to a data logger acquisition center (e-PépiPIAF 1.1, Hydrasol, Le Plessis Robisson, France). Straight and unbranched segments of main branches longer than 5 *cm* were selected for the installation of LVDT dendrometers, which were mounted using a custom-made stainless steel Invar holder (alloy with minimal thermal expansion) adapted for lavender (Lamacque et al., 2020). At the end of the experiment, final branch diameter was measured with a caliper (Burg Wächter, 0.01 *mm* accuracy) at the same location as the LVDT dendrometer measurements.

Three parameters were derived from branch diameter variations: spring relative growth, autumn relative growth, and maximum daily shrinkage (MDS).

Spring relative growth (%) was calculated as:

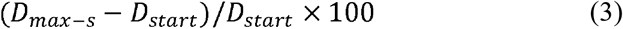

where *D_max_*_–*s*_ is the seasonal maximum diameter recorded for each individual at midnight, and *D_start_* corresponds to the diameter measured on 01/04/2025 at midnight, the date at which growth initiation was observed. No spring growth data were available for 2024, as dendrometers were installed too late in that season.

Autumn relative growth (%) was calculated as:

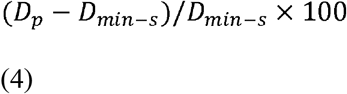

where *D_min_*_–*s*_ is the seasonal minimum diameter recorded for each individual at midnight, and *D_p_* corresponds to the diameter measured on 03/11/2024 for the 2024 season and on 15/11/2025 for the 2025 season, both selected as the time points at which the diameter signal reached a plateau, indicating the end of growth.

MDS (%) was calculated as:

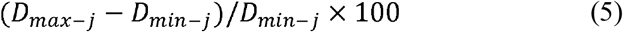

where *D_max_*_–*j*_ and *D_min_*_–*j*_ are the maximum and minimum daily diameters. MDS was calculated during periods of high evaporative demand, including heatwave events: from 2024-07-18 to 2024-07-21, from 2024-07-29 to 2024-08-02, from 2025-06-27 to 2025-07-03, and from 2025-08-08 to 2025-08-14. Additionally, MDS was calculated over longer summer heat periods, corresponding to 2024-07-23 to 2024-08-31 and 2025-07-25 to 2025-08-27. Finally, MDS was also assessed during the flowering period in 2025, which occurred between 2025-06-03 and 2025-06-26, depending on individual plants.

### 2.6. Water potentials

Water potentials (Ψ) were measured using a Scholander-type pressure chamber (PMS Instrument, Albany, OR, 565 USA) (Scholander et al., 1965). For each sampling date, equal numbers of individuals were sampled in the control and midday-shaded treatments, with one sample collected per plant; sample sizes ranged from 5 to 12 plants per treatment for lavandin, and 8 plants per treatment for parsley and valerian. Measurements were carried out during sunny conditions and after several consecutive sunny days without rain, from June to September. Samples were collected and immediately put in airtight bags under cooled conditions before measurement. Measurements were performed within a maximum of 2 h 30 *min* after sample collection. For predawn water potential (Ψ_pd_), which is commonly used as a proxy for soil water availability, samples were collected before sunrise, between 3 and 4 am (solar time), while for midday water potential (Ψ_md_), samples were harvested between 12:00 am and 1:00 pm (solar time). Measurements were carried out on one leafy branch per plant for lavandin, approximately 10 *cm* long and located on the south-facing upper part of the plant, and on one healthy leaf per plant for valerian and parsley, located on the south-facing upper part of the rosette.

### 2.7. Photochemical efficiencies

Maximal and light-adapted state photochemical efficiencies of the Photosystem II (PSII) were measured using a chlorophyll Fluorescence Monitoring System Manual according to the manufacturer instructions (FMS1+, Hansatech Instruments Ltd., UK). The maximum photochemical efficiency of PSII (*F_v_/F_m_*) was measured only in 2025 after dark adaptation of leaves using dark leaf clips supplied with the apparatus. Leaves were dark-adapted for at least 30 min prior to measurements. Measurements were performed on three dates for lavandin and four dates for parsley and valerian at solar noon. Light adapted state photochemical efficiency of PSII (ΦPSII = F’_v_/F’_m_) (Andrews et al., 1993; Baker, 2008; Nogues et al., 2001) was measured at solar noon under natural sunlight conditions on clear days and repeated at multiple time points throughout the summer. In 2024, measurements were performed on four dates for parsley and valerian. In 2025, measurements were performed on 11, 9 and 10 dates for lavandin, valerian and parsley, respectively.

For lavandin, measurements were performed on a fully expanded mature leaf from the current year’s growth, between the 4th and 7th leaf node on a south-facing shoot, fully exposed to sunlight and unshaded by upper leaves. For parsley and valerian, measurements were conducted on healthy, fully developed outer leaves on the southern side of the rosette, also fully exposed to sunlight.

For each species, measurements were conducted on 12 different plants with one leaf per plant for each treatment.

### 2.8. Biomass yields

For Lavandin, flowering stems were collected when 50% of the flowers on the spike were open (Lammerink et al., 1989; Saunier et al., 2022), i.e. on June 25, 2025. For each treatment, three sets, each composed of ten plants, were harvested and weighed immediately after harvest. For each set, 2 *kg* of fresh biomass were sampled and dried in a ventilated oven at 28 *°C* for seven days. Dry biomass was then determined per plant.

Valerian planted on June 3, 2024 was harvested on March 14, 2025, while valerian planted on May 19, 2025 was harvested on December 15, 2025. In 2024, all plants were harvested (29 plants per treatment). In 2025, 50 plants per treatment were harvested. After harvest, aboveground biomass was separated from roots. Roots were dried in a ventilated drying oven at 28 *°C* for one week. Root dry biomass was then determined per plant.

Parsley was harvested when most of plants reached 20 *cm* in height; harvest dates and growth varied according to treatment and years. In 2024, a single harvest was performed, with 10 *m* of row per treatment harvested, dried, and dry biomass yield was calculated per linear meter. In 2025, three harvests were conducted for the midday-shaded treatment and two for the control. The first harvest followed the same protocol as in 2024. For the second harvest, 3 *m* segments per row were harvested in the midday-shaded treatment, while 10 *m* per row were harvested in the control. The third harvest, conducted only the midday-shaded treatment, followed the same protocol as the second harvest. After each harvest, parsley foliar biomass was dried and weighed, and annual yield was calculated and expressed per linear meter per year.

### 2.9. Essential oil yield

Lavandin essential oil was extracted from dried stemless flowers by hydrodistillation. For each treatment, three samples were analyzed. Approximately 50 *g* of dried flower samples were placed in a Clevenger-type apparatus with 1.5 *L* of distilled water. The distillation was continued for 45 min after the first drop of essential oil was collected. The essential oil content was calculated as the ratio between the oil volume collected (*mL*) and the dry plant material weight (*g*) and expressed as v/w (*%*).

Valerian essential oil was extracted by hydrodistillation. Three samples were analyzed per treatment and each sample consisted of 40 *g* of dried and ground roots subjected to hydrodistillation in the presence of 500 *mL* of distilled water. Essential oil extraction was performed for 4 *h* using a Clevenger-type apparatus equipped with a 0,5 *ml* trap containing 1,2,4-trimethylbenzene R as the collecting solvent. The recovered essential oil was separated and the essential oil content was determined based on the extracted oil volume relative to the dry mass of plant material (v/w %).

### 2.10. Statistical analysis

Statistical analyses were performed using R software (R Core Team, version 4.5.0, 2025). Microclimatic variables and MDS were analyzed using linear mixed-effects models (LMM), with treatment considered as a fixed effect and sensor identity and measurement date included as random effects.

The normality of distribution of all other variables was assessed using the Shapiro–Wilk test. Comparisons between control and midday-shading condition were performed separately for each species using either Student’s t-test, Mann–Whitney test or general linear model (GLM), depending on data distribution. Statistical significance was determined at a threshold of p < 0.05, while trends were considered when 0.05 ≤ p < 0.10.

In the figures and tables, asterisks indicate significant differences between midday shading and control treatments (* p ≤ 0.05; ** p ≤ 0.01; *** p ≤ 0.001) and the points indicate a marginal trend between midday shading and control treatments (*p <* 0.1).

## 3. Results

### 3.1. Effect of midday-shading on air microclimate parameters

To characterize the environmental modifications induced by the midday-shading treatment, we first assessed its effects on the main microclimatic parameters, including light availability, air temperature, vapor pressure deficit (VPD), and relative humidity throughout the vegetative seasons.

Midday shading consistently reduced the PAR throughout the vegetative seasons, with the strongest reductions occurring during spring and late summer with a lower sun elevation in the sky (Fig. 2). Midday shading significantly reduced both the daily sum of PAR and the maximum instantaneous PAR throughout the vegetative season (Fig. 2). The magnitude of these reductions varied according to the season. The reduction in the daily sum of PAR ranged from 19.9% in June to 55.9% in August, with an intermediate reduction of 30.0% in July compared with control conditions. These seasonal differences coincided with variations in the daily duration of shading, which averaged approximately 10 h in April, 6 h 10 min in May, 3 h in June, 3 h 50 min in July and 7 h 30 min in August. Similarly, midday shading decreased the maximum instantaneous PAR, with reductions ranging from 15.4% in July to 45.8% in April. The smallest relative reductions were observed in June and July, when peak PAR values were highest, whereas the largest reductions occurred in April and August, when maximum PAR was lower.

**Fig. 2.**
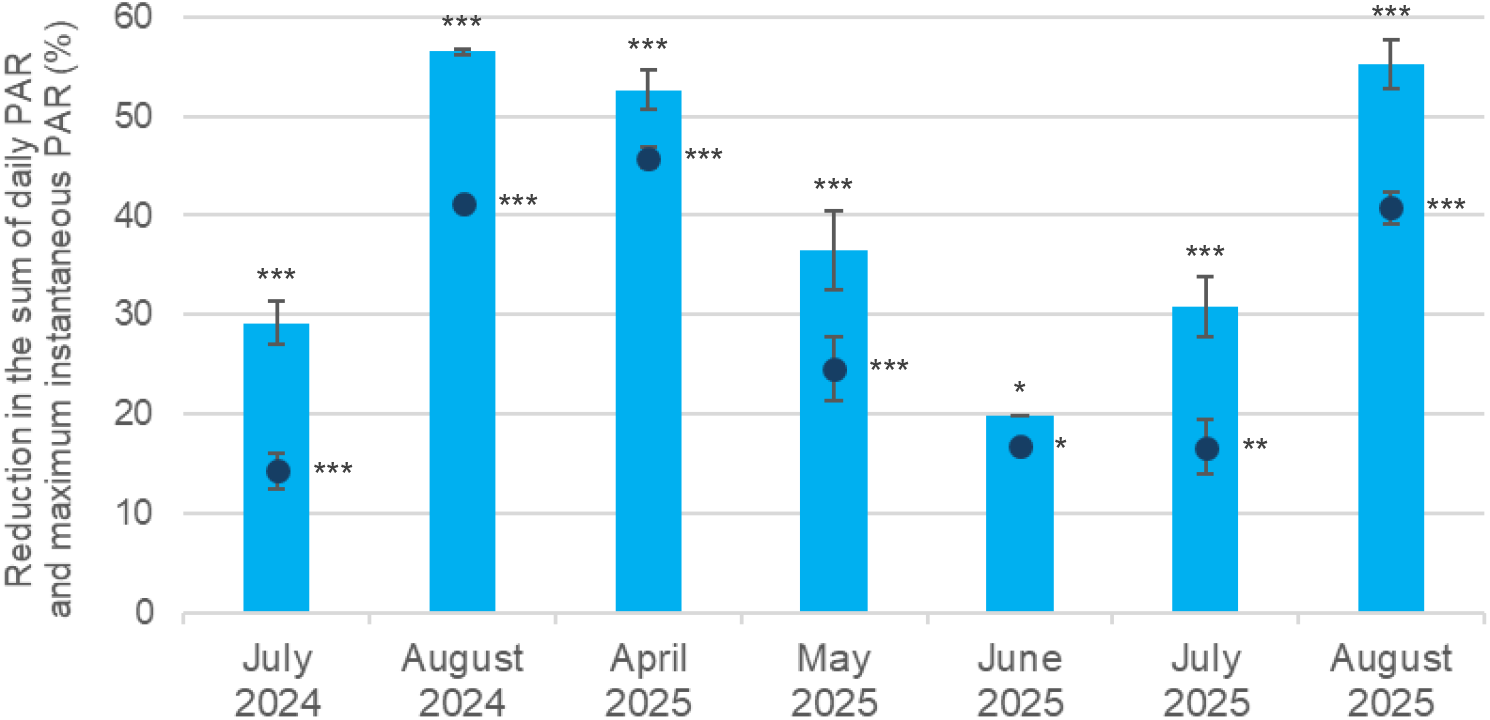
Effect of midday shading on daily cumulative and maximum instantaneous PAR. The reduction in the sum of daily PAR (bars) and maximum instantaneous PAR (points) under shaded conditions, expressed as a percentage of the values measured under control (full sunlight) conditions, was recorded during July and August 2024 and from April to August 2025. Data represents mean values from three sensors and bars indicate standard deviation (SD).

Midday shading had no statistically significant effect on maximum sheltered air temperature under most conditions (Fig. 3), neither for monthly averages (LMM, 0.106 < p < 0.391) nor for the five hottest days of 2024 (LMM, p = 0.67). However, a trend toward lower air temperatures was observed under shaded conditions, particularly for extreme heat events (Fig. 3). Indeed, midday shading seems to lower the daily maximum sheltered air temperature slightly in 2025, with an average decrease of 1.83 ± 0.96 °C (LMM, p = 0.0664).

**Fig. 3.**
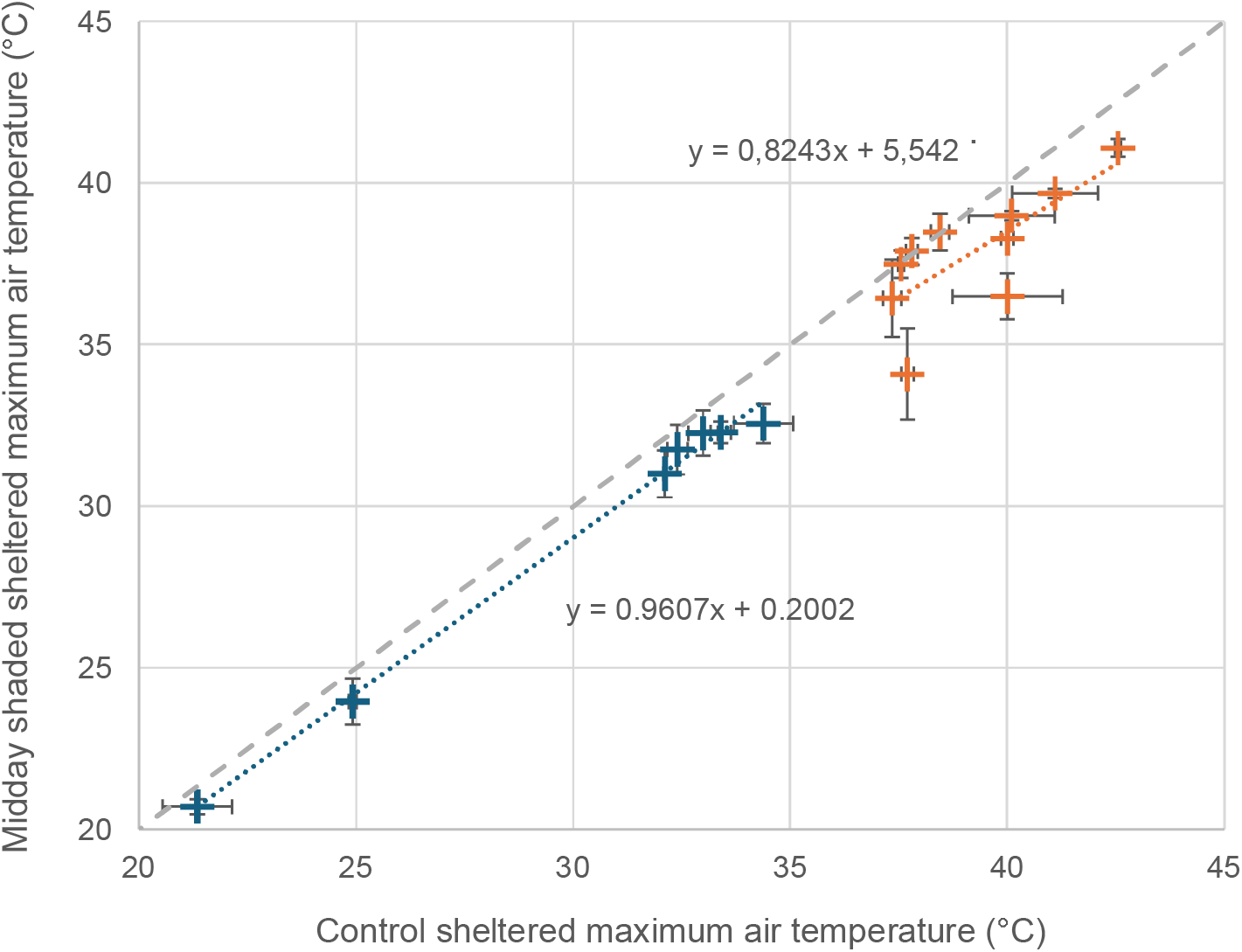
Effect of midday shading on maximum sheltered air temperature. Relationship between sheltered air temperature (*°C*) measured under control (x-axis) and under midday shading (y-axis) conditions. Blue symbols correspond to data recorded during June–July 2024 and April–August 2025, while orange symbols correspond to the five hottest days of 2024 and 2025. Values represent means from two sensors, and bars indicate the SD. The gray dashed line represents the 1:1 relationship, while the dotted lines represent the linear trend lines fitted to the corresponding datasets (blue and orange, respectively).

Midday shading had also little effect on atmospheric moisture conditions (Figure 4). Maximum VPD was generally unaffected throughout the vegetative seasons, with only a marginal reduction of 4.9% observed in June 2025 (LMM, *p* = 0.091). Similarly, VPD measured during the five hottest days was generally not affected in 2024 but showed a marginal 12.2% decrease in 2025 (LMM, *p* = 0.059).

**Fig. 4.**
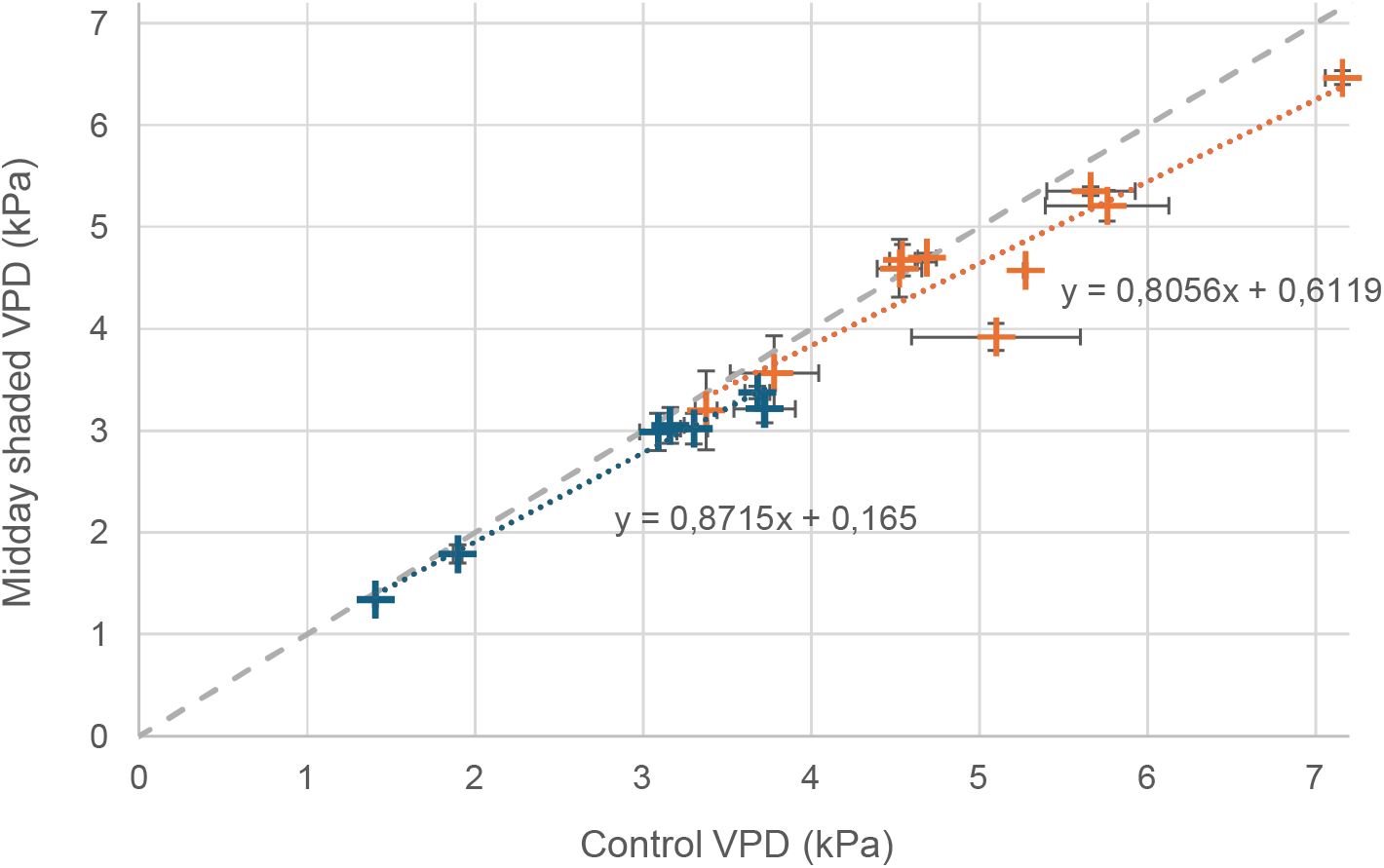
Effect of midday shading on maximum vapor pressure deficit. Relationship between maximum VPD (*kPa*) measured under control (x-axis) and under shading (y-axis) conditions. Blue symbols correspond to data recorded during June–July 2024 and April– August 2025, whereas orange symbols correspond to the five hottest days of 2024 and 2025. Values represent means from two sensors, and error bars indicate the SD. The gray dashed line represents the 1:1 relationship, while the dotted lines represent the linear trend lines fitted to the corresponding datasets (blue and orange, respectively).

No significant difference in wind speed or direction was found between the control and midday shading treatments (data not shown).

### 3.2. Integrated ecophysiological responses to midday shading

To evaluate how the radiation reduction induced by midday shading translated into plant functioning, we assessed a suite of complementary ecophysiological traits and parameters related to leaf temperature, morphology, water status, photosynthetic performance, growth dynamics and productivity throughout the vegetative season.

We first examined the effect of midday shading on leaf surface temperatures. Midday shading at solar noon significantly reduced leaf surface temperature in all three species, with a mean decrease of 6.89 °C for lavandin (GLM, p<0,00003), 5.60 °C for valerian (GLM, p = 0,00001) and 2.97 °C for parsley (GLM, p = 0,0027) compared with the control treatment (Figure 5). In the morning, prior to the midday shading period, the leaf temperature was significantly lower under the midday-shading treatment in valerian, by 1.57°C (GLM, p = 0.0013). For lavandin, a similar trend was observed, with leaf temperature being 1.49°C lower under midday shading treatment, although the difference was not statistically different (GLM, p = 0.055). For parsley, there was no difference between treatments (GLM, p = 0.424).

**Fig. 5.**
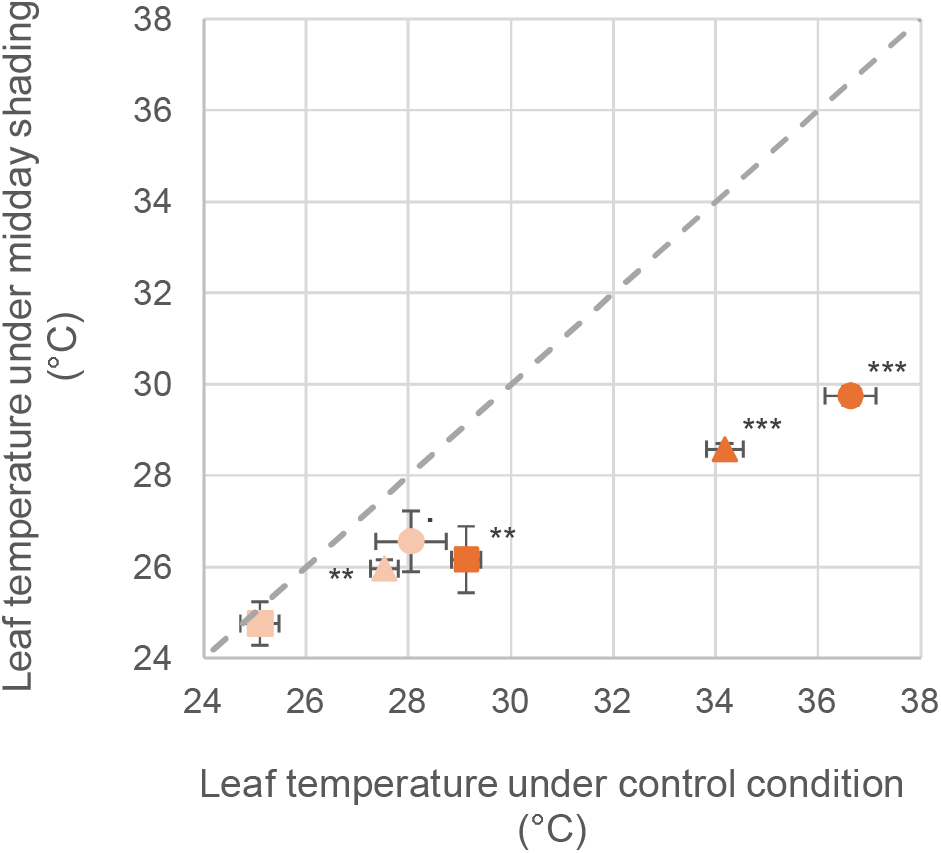
Effect of midday shading on leaf surface temperature in the three species. Relationship between leaf surface temperature (°C) measured during full sunlight (x-axis) then during shading (y-axis) periods for lavandin (circles), valerian (triangles), and parsley (squares). Light colors represent leaf surface temperature in the morning just before shading while dark colors represent surface temperature at solar noon during the shading period. Values represent means of three measurements and error bars indicate the SD. The gray dashed line represents the 1:1 relationship.

During measurements of leaf temperature in lavandin, air temperature under shelter was 31.1 °C in the control and 29.7 °C under midday shading at solar noon. During measurements of leaf temperature in parsley and in valerian, air temperature was 31.9 °C in the control and 30.5 °C under midday shading. In the morning, before the onset of shading, air temperature was 26.1 °C in the control and 26.0 °C under midday shading while measuring leaf temperature for lavandin, and 26.5 °C and 26.3 °C, respectively, for parsley and valerian.

Midday shading induced changes in morphological traits such as the LMA and mean LA in the investigated species Lavandin and Valerian (table 1). In response to midday shading, mean LA increased in both lavandin and valerian plants, while LMA decreased.

**Table 1.** Effects of midday shading on mean leaf area (LA) and leaf mass area (LMA) in lavandin and valerian. Values represent means ± SD (*n* = 12).

| Year | Species | Treatment | LA (mm<sup>2</sup>) | LMA (g.m<sup>-2</sup>) |
| --- | --- | --- | --- | --- |
| 2024 | Lavandin | Control | 83,5 ± 16,1 | 121 ± 16,4 |
|  | Lavandin | Midday shading | 106 ± 12,5 *** | 83,5 ± 13,5 *** |
| 2025 | Lavandin | Control | 82,0 ± 11,1 | 149 ± 8,81 |
|  | Lavandin | Midday shading | 102 ± 15,8 ** | 108 ± 8,92 *** |
|  | Valerian | Control | 6447 ± 2013 | 70,1 ± 7,34 |
|  | Valerian | Midday shading | 12525 ± 4406 *** | 58,5 ± 6,42 *** |

We observed that midday shading negatively affected autumn relative growth in 2024 and 2025 (Table 2). Midday shading did not significantly affect spring relative growth in 2025 (Mann–Whitney test, p = 0.222). Spring relative growth could not be assessed in 2024 because dendrometers were only operational from July 1st.

**Table 2:** Effect of midday shading on spring and autumn relative radial growths in lavandin. Relative growth (%) was calculated as the increase in branch diameter relative to the minimum branch diameter measured before the onset of growth period. Values represent means ± SD (n = 5). NA means not analyzed.

| Year | Treatment | Spring relative growth (%) | Autumn relative growth (%) |
| --- | --- | --- | --- |
| 2024 | Control | NA | 63,3 ± 14,4 |
|  | Midday shading | NA | 25,8± 10,2 ** |
| 2025 | Control | 29,2 ± 10,7 | 24,8 ± 6,51 |
|  | Midday shading | 18,9 ± 2,55 | 12,6 ± 4,72 * |

To evaluate the effect of midday shadding on soil and plant water status, we measured predawn (Ψ_pd_) and midday (Ψ_md_) water potentials (Figure 6), during periods expected to be water-limited in summer (see materials and methods). The effects of midday shading on shoot water potentials varied among measurement dates but overall tended to increase Ψ_md_ and, to a lesser extent, Ψ_pd_. In contrast, no treatment effect was detected in parsley and valerian because regular irrigation maintained Ψ values above −0.3 MPa throughout the experiment, beyond the operating range of the pressure chamber.

**Fig. 6:**
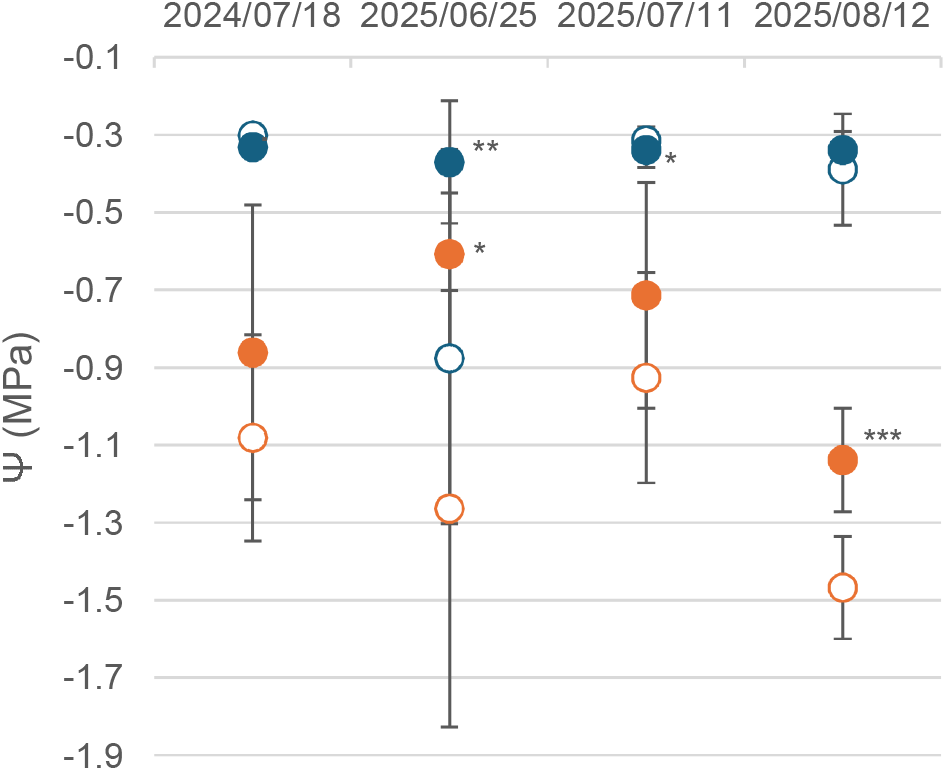
Effect of midday shading on shoot water potentials in lavandin. Predawn (Ψ_pd_, blue symbols) and midday (Ψ_md_, orange symbols) water potentials were measured under control (open circle) and midday shading (filled circle) conditions at four dates during the summer periods. Values represent means of five to twelve measurements and error bars indicate the SD.

Midday shading improved both light-adapted (F’_v_/F’_m_) and maximal (F_v_/F_m_) photochemical efficiency in the three MAP species (Figure 7). Improvements in (F’_v_/F’_m_) were consistent across the experiment, whereas shading-induced increase in F_v_/F_m_ were more variable, being more pronounced in lavandin and valerian than in parsley.

**Fig. 7.**
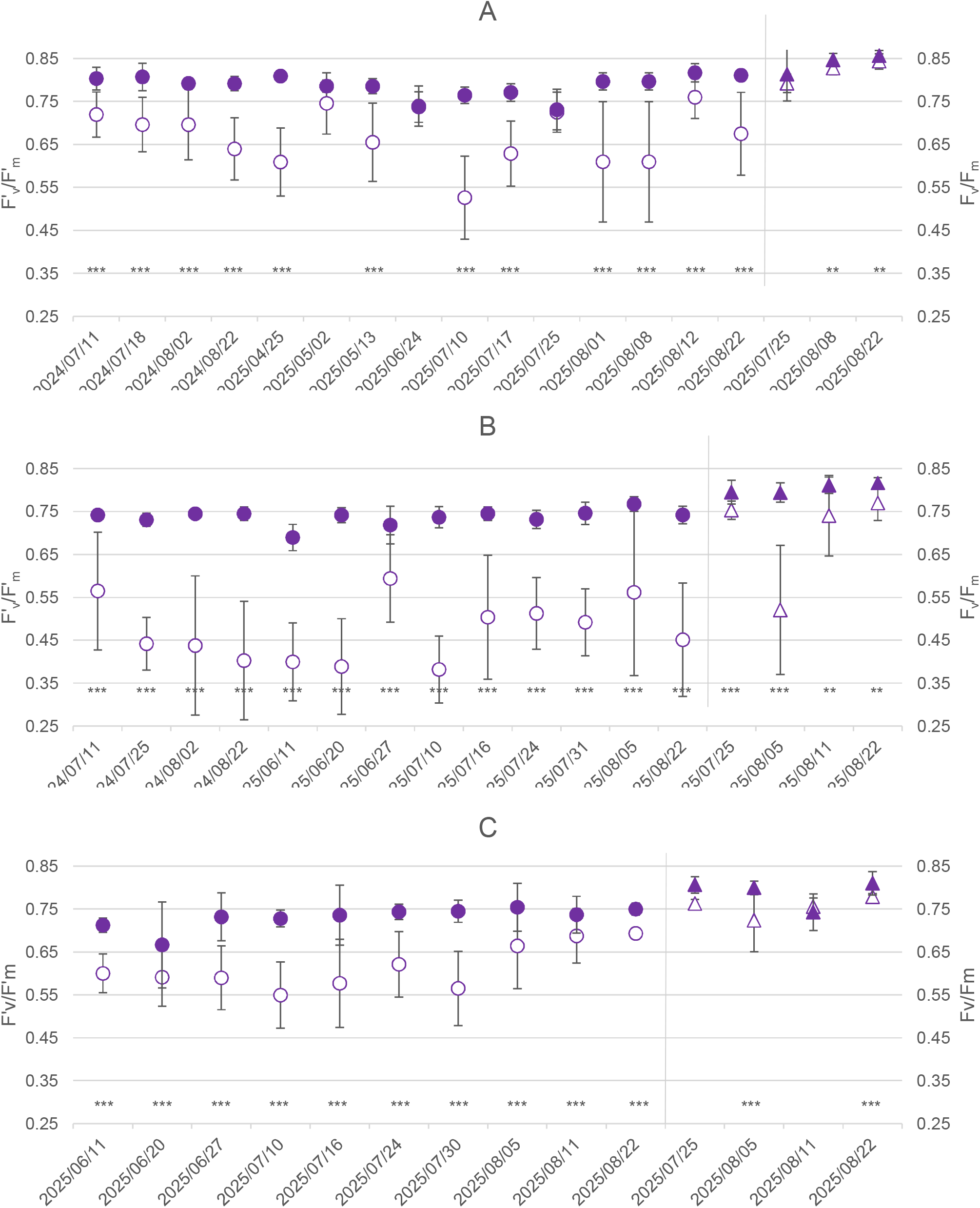
Effect of midday shading on the light-adapted and maximal photochemical efficiencies in the three species. Light-adapted F’_v_/F’_m_, left side of the graph, circles), and maximal F_v_/F_m_, right side of the graph, triangles) photochemical efficiencies were measured in lavandin (A), valerian (B), and parsley (C). Open symbols represent control plants, whereas filled symbols correspond to midday-shaded plants. Values represent means (n=12) and error bars indicate the SD.

Midday shading reduced maximum daily branch shrinkage (MDS) in lavandin during the heat periods of 2024 (Table 3). Significant reductions ranging from 32.8% to 39.7% were observed in 2024 during all heatwave events and over the extended summer heat period (LMM, *p* < 0.025), whereas no significant effect was observed on MDS (LMMs, *p* > 0.362) in 2025.

**Table 3:**
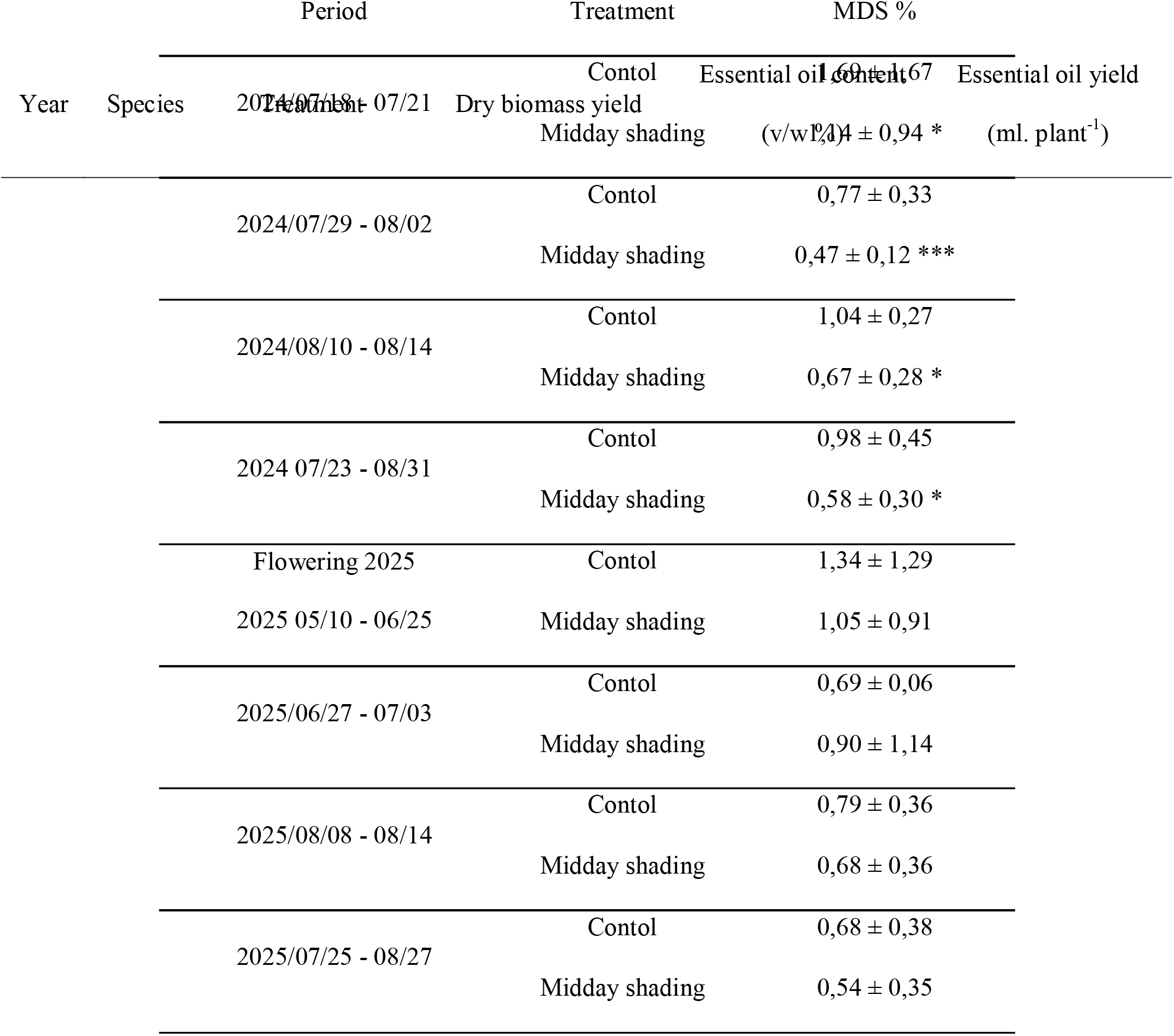
Effect of midday-shading on maximum daily shrinkage of branch in lavandin during different periods. The Maximum daily shrinkage (MDS, %) of lavandin branch was scored during different periods correspond to flowering (from 2025/06/03 to 2025/06/26, depending on individual plants), and to heat periods indicated by date ranges. Values represent means ± SD (n = 5). The calculation formula is provided in the “Materials and Methods” section.

Midday shading significantly affected biomass production, with contrasting responses among species (Table 4). Floral biomass decreased in lavandin. Root biomass increased in valerian under midday shading treatment in 2024 but not in 2025. In contrast, essential oil content remained unaffected by midday shading treatment in both lavandin and valerian. Consequently, essential oil yield largely followed biomass production, decreasing in lavandin in 2025 and increasing in valerian in 2024, while no significant differences were observed in the other harvests. For parsley, midday shading increased dry biomass yield by 75% in 2024 and 51% in 2025. In 2024, shaded plants were harvested in October, whereas control plants reached the 20-cm harvest threshold only in late November. In 2025, the first two harvests occurred earlier under midday shading (1 August and 9 September) than in the control (14 August and 26 September), and shaded plants allowed an additional third harvest on 27 October, while control plants did not reach the harvest threshold again.

**Table 4:**
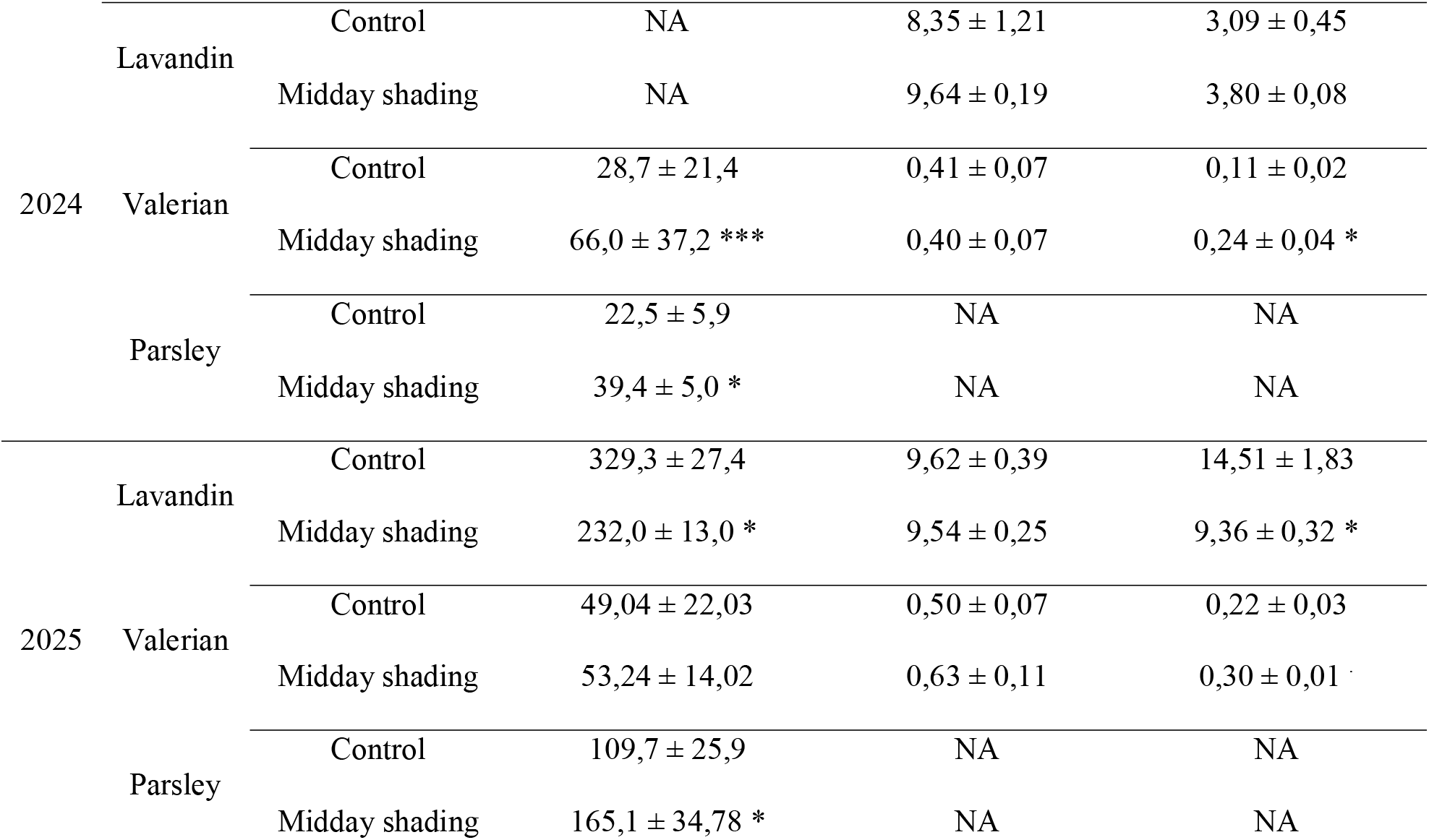
Effect of midday-shading on yields in the three species. Dry biomass and essential oil yields were measured in lavandin, valerian and parsley grown under control and midday shading treatments in 2024 and 2025. Harvested organs corresponded to floral stems for lavandin, roots for valerian, and leaves for parsley. Biomass yields are expressed as *g.plant ¹* for valerian and lavandin and *g.linear m ¹.year ¹* for parsley. Essential oil content *(% v/w*) and essential oil yield (*ml.plant ¹*) were also measured for lavandin and valerian.

## 4. Discussion

### 4.1. Midday shading primarily modifies the radiative environment rather than the crop air microclimate

Although the vertical shade nets substantially modified the radiative environment perceived by the crops, their effects on the surrounding microclimate remained limited. Seasonal differences in PAR attenuation reflected changes in solar elevation, with greater reductions in spring and late summer due to the longer daily duration of shading. This temporal pattern differs from the heterogeneous light environments encountered in agroforestry systems, where light availability depends on multiple interacting factors including tree age and size, canopy architecture, planting density, row orientation, latitude, season and time of day (Dupraz et al., 2018; Feldhake and Belesky, 2009; Pardhi et al., 2020; Peng et al., 2025). Agrivoltaic systems also generate heterogeneous light environments, with irradiance reductions ranging from 30 to more than 80% depending on panel configuration, and daily shading varying between 4 and 88% (Disciglio et al., 2025; Juillion et al., 2022; Pang et al., 2019). The overall PAR reduction measured in this study was generally lower than under dense tree canopies, some agrivoltaic systems or permanent shade nets, its temporal distribution more closely resembled the dynamic light environment of open agroforestry systems (Mantino et al., 2021).

The limited effects on air temperature and VPD are consistent with modest reductions in VPD (3–4%) reported under permanent 50% shade nets (Montanaro et al., 2009). Others studies showed contrasting effects on air temperature, ranging from no effect to decreases of 1–3 °C depending on shading intensity, net characteristics and local climatic conditions (Mupambi et al., 2018). By contrast, agroforestry and agrivoltaic systems substantially modify the atmospheric microclimate. In agroforestry systems, daytime air temperature could be decreased by 1 to 2 °C, with cooling effects reaching up to 6 °C during extreme heat events, while simultaneously reducing canopy temperature (Gosme et al., 2016; Inurreta-Aguirre et al., 2018; Kanzler et al., 2019; Karvatte et al., 2020; Sida et al., 2018). Unlike shade nets, trees also increase atmospheric humidity through transpiration, with reported increases in relative humidity ranging from 3 to 12% depending on the agroforestry system (Campi et al., 2009; López-Díaz et al., 2026; Von Arx et al., 2012). Agrivoltaic systems similarly reduce air temperature up to 4°C (Ali Abaker Omer et al., 2025; Magarelli et al., 2024; Marrou et al., 2013; Pang et al., 2019), increase relative humidity around 10-14% (Barron-Gafford et al., 2019; Juillion et al., 2022) and lower VPD (Marrou et al., 2013), although responses vary with panel orientation and configuration and local climate. These stronger responses arise because trees combine radiative shading with evapotranspirative cooling (Blanchet et al., 2022), whereas photovoltaic panels additionally create a more spatially homogeneous radiative environment over the cropped area (Adolfo et al., 2023). Moreover, windy conditions during the experiment probably further reduced these effects by enhancing air mixing within the canopy, consistent with Disciglio et al. (2025), who reported only modest reductions in air temperature (0.3 - 0.5 °C) under similarly windy conditions.

The primary effect of the vertical shade nets was to reduce the radiative load absorbed by the leaves, reducing the leaf surface temperature in all three species at solar noon (Figure 5). Consistently, leaf temperature under full sunlight exceeded ambient air temperature by approximately 5.56 °C in lavandin and 2.33 °C in valerian, indicating a strong radiative heating of the foliage. By contrast, parsley leaf temperature remained approximately 2.72 °C below air temperature even in the control treatment at solar noon, suggesting a fundamentally different leaf energy balance or a better transpiration efficiency. Under midday shading at solar noon, lavandin leaf surface temperature became almost identical to the sheltered air temperature, exceeding it by only 0.07 °C. In valerian and parsley, leaf surface temperature remained below sheltered air temperature, by 1.90 °C and 4.32 °C, respectively. Unlike lavandin, both valerian and parsley were irrigated throughout the experiment, allowing sustained transpiration and evaporative cooling, which likely contributed to their lower leaf temperatures (Farella et al., 2022; Jones and Rotenberg, 2011). This interpretation is further supported by the consistently high Ψ_md_ measured in both species (Ψ_md_ > −0.3 MPa), indicating that plants remained well hydrated despite the high evaporative demand. In parsley, this effect was probably reinforced by its finely dissected leaves, which intercept less solar radiation and reduce boundary layer thickness, thereby enhancing convective and transpirational cooling (Leigh et al., 2017; Vogel, 2009). Consequently, parsley maintained leaf temperatures below ambient air temperature even under full sunlight. The effect of leaf cooling observed here is consistent with previous studies conducted on other species under agrivoltaic systems, shade nets and agroforestry, which generally reported larger reductions in leaf or canopy temperature (2–9 °C) than in air temperature (Disciglio et al., 2025; Karvatte et al., 2020; Lott et al., 2009; Marrou et al., 2013). Interestingly, leaf surface temperature was already lower in the midday-shading treatment before shading was applied, particularly in valerian and, to a lesser extent, in parsley. This observation supports the hypothesis that, in these well irrigated species, sustained transpiration contributed to leaf cooling throughout the day.

The reduction in leaf temperature induced by midday shading effectively alleviated daytime water stress by reducing transpirational demand. This was reflected in the consistently higher Ψ*_md_* observed under shading conditions in lavandin throughout the experiment, with significant differences recorded during the periods of greatest atmospheric demand. This indicates that the limiting radiative loading and leaf overheating enabled plants to maintain a more favorable water status. As mentioned above, predawn water potential was significantly higher under midday shading only on 25 June 2025. This measurement was performed during the flowering period, when plant water requirements are particularly high (Wang and Xu, 2026), and coincided with a heatwave. Moreover, no substantial rainfall had occurred since 4 and 5 June 2025, when a cumulative 15 mm of precipitation was recorded. By reducing daytime transpiration during the heatwave, shaded plants likely extracted less water from the soil, allowing a more complete overnight rehydration. A similar mechanism was reported by Mira-García et al. (2022), who showed that shading slowed soil water depletion and delayed the onset of severe water stress, although differences disappeared once soil water reserves became largely exhausted.

Midday shading had no influence of Ψ*_pd_, likely* because of the high irrigation, with Ψ*_pd_*, remaining similar between treatments on most measurement dates. Because Ψ*_pd_*was measured after nocturnal hydraulic equilibration between soil and plants, it primarily reflects soil water availability explored by roots and the capacity of plants to fully rehydrate overnight rather than the atmospheric conditions experienced during the previous day (Kangur et al., 2017). The only improvement due to partial shading occurred on 25 June 2025, when Ψ*_pd_* increased by approximately 0.5 MPa under midday shading conditions. The physiological significance of this isolated response is discussed later in relation to the effects of midday shading on leaf temperature and midday water potential.

Overall, the influence of midday shading became more pronounced under warmer conditions, suggesting that its primary effect was to buffer short periods of intense thermal and evaporative demand rather than to modify the daily crop microclimate. Because leaf surface temperature and water potential were measured only during the warmest and sunniest periods, when radiative and thermal stresses were greatest and the effects of midday shading were expected to be maximal, the responses reported here likely represent the maximum benefits provided by the treatment. Under these conditions, the marked reduction in leaf temperature and the associated improvement in plant water status, despite only minor changes in ambient air conditions, highlight leaf temperature as an integrative indicator of the stress experienced by plants. Conversely, during the majority of days with lower radiation and evaporative demand, the effects of midday shading were likely much weaker. The extent to which these changes translated into morphological and physiological responses was examined below.

### 4.2. A midday reduction of radiation was sufficient to modify plant functioning and development

In the present study, both lavandin and valerian exhibited a lower LMA together with an increase in mean LA under midday shading, indicating that even this partial and temporary reduction in irradiance was sufficient to trigger morphological acclimation to reduced light availability. Midday shading also affected secondary growth in lavandin, as shown by the significant reduction in autumn radial growth, and to a lesser extent in spring radial growth. Radial growth is known to depend on light availability, as increasing shade can reduce photosynthetic carbon assimilation, thereby decreasing cambial activity and ultimately limiting secondary growth. (Greis and Kellomäki, 1981; Holtta et al., 2010; Lopes et al., 2015; Yasuda et al., 2018). Together, these results indicate that temporary shading around solar noon were sufficient to induce persistent changes in plant development. They further suggest that the effects of midday shading cannot be explained only by the reduction in leaf temperature, but also reflect the consequences of reduced radiation availability, which were particularly pronounced during spring and from August onwards.

The physiological measurements were intentionally performed during the periods expected to impose the greatest thermal and evaporative constraints, because the objective was to determine whether midday shading protects plants when environmental stress is maximal. In this context, the reduction in leaf surface temperature observed under midday shading likely represents the primary mechanism underlying the subsequent physiological responses. By limiting leaf overheating during the highest irradiance, shading reduced the intensity of thermal and radiative stress experienced by the photosynthetic apparatus. This protective effect was consistently reflected in higher F’_v_/F’_m_ and F_v_/F_m_ values under shading conditions. The increase in F’_v_/F’_m_, measured under ambient light at solar noon, indicates that PSII maintained a higher operating photochemical efficiency during periods of maximum thermal and radiative stress, suggesting that midday shading alleviated the immediate inhibition of photosynthesis caused by excess irradiance (Baker, 2008; Lu and Zhang, 1999). Similarly, the higher F_v_/F_m_ values measured after dark adaptation indicate that the photosynthetic apparatus recovered more completely from daytime stress, reflecting a lower degree of sustained photoinhibition and a better preservation of PSII integrity. Since F_v_/F_m_ values close to 0.80 are generally associated with a non-stressed photosynthetic apparatus, whereas lower values indicate photoinhibition or PSII damage (Herbette et al., 2005; Jing et al., 2009; Montanaro et al., 2009), ours results suggest that midday shading protected PSII from both the immediate and the lasting effects. Primarily used to assess photoinhibition, both F’_v_/F’_m_ and F_v_/F_m_ can also be disturbed by water deficit through stomatal closure (Lu and Zhang, 1999). However, parsley and valerian were maintained under well-irrigated conditions throughout the experiment, with both Ψ*_pd_* and Ψ*_md_*consistently above - 0.3 MPa, indicating no water stress. The higher F’_v_/F’_m_ and F_v_/F_m_ values under midday shading therefore primarily reflect the alleviation of thermal and radiative stress rather than changes in plant water status. In lavandin, the improved chlorophyll fluorescence parameters observed under midday shading likely also reflect the alleviation of radiative and thermal stress. Indeed, we assume that photochemical efficiency in Lavandin was not substantially affected by water stress, as the stomatal closure likely did not occur. The Ψ*_md_* measured in our study remained higher than the water potential reported to induce stomatal closure in this species (Lamacque et al., 2020). However, midday shading significantly reduced MDS during the 2024 heatwaves, indicating lower branch dehydration and reduced daily water stress when compared to control plants. The weaker difference for MDS in 2025 may reflect the greater establishment of lavandin plants during their second vegetative season, making them less sensitive to mild water deficits.

Together, these results indicate that even though midday shading produces only minor changes in the surrounding microclimate, it efficiently reduces the intensity of the most stressful conditions experienced by plants during hot sunny days.

### 4.3. Physiological improvements do not necessarily translate into higher yields

Although midday shading improved the physiological status of the three species by alleviating thermal and radiative stress, these benefits did not translate into similar agronomic responses. Indeed, it induced contrasting biomass responses among species, decreasing lavandin floral biomass while substantially increasing parsley shoot biomass and valerian root biomass. Lavandin floral biomass decreased by approximately 26%, whereas parsley shoot biomass increased by 51 to 75% and valerian root biomass has more than doubled in 2024. These contrasting responses likely reflect a trade-off related to PAR between the alleviation of thermal, radiative and water stress and the reduction in carbon assimilation and biomass production. In lavandin, for which reproductive shoots constitute the harvested product, the lower PAR availability may have constrained floral biomass production despite the improved physiological status of shaded plants under stressful conditions. In *Lavandula dentata*, for example, 50% shading reduced flower yield by approximately 37% and essential oil yield by 32 to 47% (Mambrí et al., 2018). In contrast, shade acclimation commonly involves increased biomass allocation to leaves to maximize light interception under reduced irradiance (Poorter et al., 2012). Because leaves constitute the harvested organ in parsley, such adjustments may have contributed to the positive biomass response observed under midday shading. Parsley also benefited from midday shading also through faster attainment of the 20-cm height harvest threshold. In 2024, shaded plants were harvested approximately 1.5 months earlier than control plants. In 2025, the first two harvests also occurred earlier under shade, and a third harvest was possible in late October only for shaded plants. Thus, for parsley, the benefits associated with stress alleviation appeared to outweigh the reduction in daily radiation. A similar trade-off may have influenced the response of valerian. Although shade acclimation generally tends to favor biomass allocation to aboveground, light-capturing organs rather than roots (Poorter et al., 2012), shading does not necessarily reduce root production. For instance (Chichaghare et al., 2026) reported no significant effect of shading on root biomass in chickpea. In the present study, valerian root biomass more than doubled under midday shading in 2024. This suggests that, under the conditions experienced that year, the benefits of reduced thermal, radiative and water stress may have compensated for the limitation in carbon assimilation associated with reduced PAR, resulting in greater root biomass. However, this positive response was not observed in 2025, highlighting the importance of environmental conditions in determining the balance between stress alleviation and limitation in carbon assimilation.

Essential oil concentration in lavandin and valerian was unaffected in the present study. Yet, numerous studies showed that shading and water stress can modify the production and composition of essential oils in aromatic species, thereby influencing product quality in addition to biomass production (Figueiredo et al., 2008; Katsoulis et al., 2022; Lalević et al., 2023; Mambrí et al., 2018; Moradkhani et al., 2010; Zubay et al., 2021). Such discrepancy between our finding and the literature may be related to the temporary shading in our study which would have limited effect on these metabolisms.

### 4.4. Species ecology would determine the balance between stress alleviation and carbon limitation

Although midday shading improved physiological functioning in all three species, the extent of the effects differ between species, as indicated by the results on leaf surface temperature (Fig. 5) and level of photochemical efficiency (Fig. 7). Moreover, its agronomic consequences differed markedly according to species ecology. Midday shading reduced floral biomass and essential oil yield in Lavandin, whereas parsley and valerian exhibited positive shoot and root biomass responses, emphasizing the strong species-specific responses of MAP Yields to shading.

Lavandin is a Mediterranean species naturally adapted to high irradiance, elevated temperatures and recurrent summer drought (Lis-Balchin, 2002; Pokajewicz et al., 2023). Under the climatic conditions of the present study, it was therefore cultivated under favorable conditions. Consequently, although midday shading consistently improved PSII functioning, the physiological benefits remained relatively limited. Dark-adapted F_v_/F_m_ values remained close to the optimal value of approximately 0.83 reported for non-stressed leaves indicating little or no sustained photoinhibition (Bjorkman and Demmig, 1987). Similarly, F’_v_/F’_m_ values remained high under midday shading (around 0.79), whereas they decreased to approximately 0.67 under full sunlight, indicating only a moderate and reversible reduction in PSII operating efficiency. Overall, the benefits of stress alleviation would be insufficient to compensate for the reduction in incident PAR, which limited carbon acquisition and ultimately resulted in lower floral biomass production and reduced radial growth. Yield reductions induced by shading have also been reported in many Mediterranean MAP species, with lavender, lemon thyme, common thyme and rosemary grown beneath photovoltaic panels showing reduced growth and lower fresh aboveground biomass (Disciglio et al., 2025).

In contrast, valerian is naturally associated with cool and moist habitats (Lee et al., 1996; Patočka and Jakl, 2010) and was cultivated under climatic conditions considerably warmer than its ecological optimum. Under these conditions, thermal and radiative stresses were likely major constraints to growth despite adequate water supply (Ψ*_pd_* and Ψ*_md_* > −0.3MPa). This interpretation is supported by the strongest response of photochemical efficiencies to midday shading in valerian compared to the two other species. Under full sunlight, F’_v_/F’_m_ values decreased to approximately 0.47, indicating a marked reduction in the operating efficiency of PSII under peak irradiance, while F_v_/F_m_ values around 0.70 reflected damage to PSII (Maxwell and Johnson, 2000). The reduction in leaf temperature and the improved preservation of PSII function under midday shading translated into a marked increase in root biomass, particularly during the first growing season. For this species, the physiological benefits of reducing thermal and radiative stress would outweigh the costs associated with reduced PAR.

Parsley exhibited an intermediate response consistent with its ecology as a temperate species. Although less sensitive to heat and drought than valerian, curly parsley is not as well adapted to intense summer radiation and high temperatures as Mediterranean species such as lavandin. This intermediate response was also reflected in photochemical efficiency (Fig. 7). F_v_/F_m_ values remained relatively high under both treatments (approximately 0.76 in the control and 0.78 under midday shading), indicating only limited sustained photoinhibition. F’_v_/F’_m_ values were close to those measured in lavandin, although consistently slightly lower. averaging approximately 0.61 under full sunlight and 0.73 under midday shading, compared with 0.67 and 0.79 in lavandin, respectively. These results indicate that parsley experienced moderate thermal and radiative stress, sufficient for midday shading to improve PSII functioning while maintaining adequate daily radiation for carbon acquisition. Consequently, the benefits of stress alleviation would outweigh the reduction in PAR, resulting in a net increase in shoot biomass in both experimental years.

Moreover, yields results observed in this study agree with previous studies showing that Mediterranean aromatic species generally exhibit reduced biomass under strong shading, whereas temperate species often tolerate or even benefit from moderate reductions in irradiance (Disciglio et al., 2025; Mambrí et al., 2018; Zubay et al., 2021). Similarly, several shade-tolerant MAPs, including yarrow, marigold, caraway, and Moldavian dragonhead, increased their yields under 30% shading. However, under stronger shading (50%), yield reductions were observed, highlighting the importance of shading intensity in determining crop responses. (Zubay et al., 2021b).

Overall, these findings demonstrate that the agronomic response to partial midday shading is primarily determined by species ecology, and would be related to the balance between the benefits of stress alleviation and the costs associated with reduced PAR. Under our experimental conditions, this balance remained unfavourable for heliophilous Mediterranean species such as lavandin. In contrast, species originating from cooler or temperate environments benefited more from midday shading. However, this balance is likely to shift as climate change increases the frequency and intensity of heatwaves associated to drought. For Mediterranean heliophilous species, the reduction in thermal and water stress could progressively outweigh the costs associated with reduced PAR, under warmer conditions.

### 4.5. Is midday shading a promising strategy for commercial production ?

Although agroforestry provides multiple ecosystem services beyond shading, including biodiversity conservation, evaporative cooling and hydraulic redistribution, it also introduces belowground competition for water and nutrients (Arenas-Corraliza et al., 2018; Artru et al., 2017; Fernández et al., 2008; Inurreta-Aguirre et al., 2018; Kanzler et al., 2019; Livesley et al., 2004; Mantino et al., 2021; Quinkenstein et al., 2009; Temani et al., 2021) and requires substantial changes in farm organization, equipment and technical expertise. Agrivoltaic systems provide another promising approach, although crop responses largely depend on panel orientation, height, spacing and mobility. Because photovoltaic systems are generally designed to maximize electricity production, light reduction may become excessive for many medicinal and aromatic plants (Disciglio et al., 2025; Juillion et al., 2022; Pang et al., 2019). Permanent horizontal shade nets, already widely used in fruit production (El-Zawily et al., 2024; Zha et al., 2022), constitue an alternative solution.

In this context, the midday shading system evaluated in the present study represents an interesting complementary approach. By restricting shading to the hours of highest thermal and radiative demand, it alleviated stress while maintaining a relatively high daily light availability. Moreover, such systems could be improved by deploying them only during the period of the year when heat and radiative stress are most likely to occur (e.g. June to August), thereby maximizing radiation interception by the vegetation during the remainder of the growing season. Their orientation and deployment period could also be adjusted according to species requirements or climatic conditions, providing a level of flexibility that is difficult to achieve with agroforestry or most agrivoltaic systems. For heliophilous species such as lavandin, shading could even be deployed only after harvest to protect plants from late-summer heatwaves and drought, when floral yield is no longer affected but high thermal and evaporative demand may impair plant recovery and reserve accumulation for the following season.

Although these results put in light the physiological and agronomic potential of midday shading, its practical implementation now requires validation under commercial production conditions. Future studies should assess not only crop productivity and quality, but also installation costs, mechanization, labour requirements and long-term economic performance. Such evaluations will determine whether temporary midday shading can become a practical climate adaptation strategy for medicinal and aromatic plants and other high-value crops.

## 5. Conclusion

This study demonstrates that midday shading primarily modify the radiative environment experienced by plants rather than the surrounding microclimate. By reducing incident PAR during the most stressful period of the day, midday shading markedly lowered leaf temperature and alleviated thermal and radiative stress, resulting in improved physiological functioning in the three MAP species. However, these physiological benefits under stressful conditions did not systematically translate into higher yields. Agronomic responses would depend on the balance between stress alleviation and the reduction in PAR for carbon acquisition, with this balance strongly influenced by species ecology and proximity to their need in climatic conditions need. In our experimental conditions, parsley and valerian, which were exposed to warmer conditions than their favorable growing environments, benefited most from midday shading, whereas lavandin, a heliophilous Mediterranean species cultivated within its current production area, maintained improved physiological functioning but showed reduced floral biomass. Overall, these findings highlight the potential of targeted midday shading as a climate-adaptation lever for MAP production, while emphasizing that its implementation should be tailored to species requirements, harvested organs and climatic conditions.

## Author contributions

**Ombeline Decuy :** Methodology, Formal analysis, Writing – original draft, Investigation. **Benjamin Lemaire :** Funding acquisition, Supervision. **Cyril Bozonnet :** Software, Data curation. **Soline Morel :** Investigation. **Clara Paredes :** Investigation. **Laurent Barroux:** Investigation. **Pascal Walser:** Investigation. **Philippe Balandier:** Conceptualization, Supervision, Methodology, Writing – review and editing, Validation. **Stephane Herbette:** Conceptualization, Funding acquisition, Supervision, Methodology, Writing – review and editing, Validation.

## Acknowledgements

The authors thank Yash Dasputre (Iteipmai) for his valuable assistance with the various measurements conducted during this study and also Laetitia Coudert (Universite Clermont Auvergne), Florence Delepine and Céline Fabien (iteipmai) for their administrative support.

## Bibliography

Adolfo, R., Kyle, P., Azad, D., Maggie, G., Serkan, A., Kirschten, H., Chad, H., 2023. Agroforestry vs. Agrivoltaic: spectral composition of transmitted radiation and implications for understory crops. 10.21203/rs.3.rs-2911844/v1

Agyare, C., Agana, T., Boakye, Y., Apenteng, J., 2017. Petroselinum crispum: A Review, in: Medicinal Spices and Vegetables from Africa: Therapeutic Potential Against Metabolic, Inflammatory, Infectious and Systemic Diseases. pp. 527–547. 10.1016/B978-0-12-809286-6.00025-X

Alam, B., Singh, R., Uthappa, A.R., Chaturvedi, M., Et., A., 2018. Different genotypes of Dalbergia sissoo trees modified microclimate dynamics differently on understory crop cowpea (Vigna unguiculata) as assessed through ecophysiological and spectral traits in agroforestry system. Agricultural and Forest Meteorology. 10.1016/j.agrformet.2017.11.031

Ali Abaker Omer, A., Li, M., Zhang, F., Hassaan, M.M.E., El Kolaly, W., Zhang, X., Lan, H., Liu, J., Liu, W., 2025. Impacts of agrivoltaic systems on microclimate, water use efficiency, and crop yield: A systematic review. Renewable and Sustainable Energy Reviews 221, 115930. 10.1016/j.rser.2025.115930

Andrews, J.R., Bredenkamp, G.J., Baker, N.R., 1993. Evaluation of the role of State transitions in determining the efficiency of light utilisation for CO2 assimilation in leaves. Photosynthesis Research. 10.1007/BF00015057

Arenas-Corraliza, M.G., López-Díaz, M.L., Moreno, G., 2018. Winter cereal production in a Mediterranean silvoarable walnut system in the face of climate change. Agriculture, Ecosystems & Environment 264, 111–118. 10.1016/j.agee.2018.05.024

Arenas-Corraliza, M.G., Rolo, V., López-Díaz, M.L., Moreno, G., 2019. Wheat and barley can increase grain yield in shade through acclimation of physiological and morphological traits in Mediterranean conditions. Sci Rep 9, 9547. 10.1038/s41598-019-46027-9

Artru, S., Garré, S., Dupraz, C., Hiel, M.-P., Blitz-Frayret, C., Lassois, L., 2017. Impact of spatio-temporal shade dynamics on wheat growth and yield, perspectives for temperate agroforestry. European Journal of Agronomy 82, 60–70. 10.1016/j.eja.2016.10.004

Baker, N.R., 2008. Chlorophyll Fluorescence: A Probe of Photosynthesis In Vivo. Annu. Rev. Plant Biol. 59, 89–113. 10.1146/annurev.arplant.59.032607.092759

Barron-Gafford, G.A., Pavao-Zuckerman, M.A., Minor, R.L., Sutter, L.F., Barnett-Moreno, I., Blackett, D.T., Thompson, M., Dimond, K., Gerlak, A.K., Nabhan, G.P., Macknick, J.E., 2019. Agrivoltaics provide mutual benefits across the food–energy–water nexus in drylands. Nat Sustain 2, 848–855. 10.1038/s41893-019-0364-5

Bernal-Basurco, C., 2026. Regulated deficit irrigation based on plant water status and Agrivoltaic systems as possible improvements on water resources management in tomato.

Betts, A.K., 2009. Land Surface Atmosphere Coupling in Observations and Models. J Adv Model Earth Syst 1, JAMES.2009.1.4. 10.3894/JAMES.2009.1.4

Bjorkman, O., Demmig, B., 1987. Photon yield of O2 evolution and chlorophyll fluorescence characteristics at 77 K among vascular plants of diverse origins. Planta. 10.1007/BF00402983

Blanchet, G., Barkaoui, K., Bradley, M., Dupraz, C., Gosme, M., 2022. Interactions between drought and shade on the productivity of winter pea grown in a 25-year-old walnut-based alley cropping system. Journal of Agronomy and Crop Science n/a. 10.1111/jac.12488

Bray, E.A., Bailey-Serres, J., Weretilnyk, E., 2000. Responses to abiotic stresses, in: American Society of Plant Biologists. John Wiley & Sons, pp. 1158–1249.

Brengi, S.H., N. Nasef, I., 2023. Alleviating the Effects of High-Temperature Stress on Parsley Plants by Foliar Application of Proline, Glycine Betaine, and Salicylic Acid. Alexandria Science Exchange Journal. 10.21608/asejaiqjsae.2023.326581

Calvin, K., Dasgupta, D., Krinner, G., Mukherji, A., Thorne, P.W., Trisos, C., Romero, J., Aldunce, P., Barrett, K., Blanco, G., Cheung, W.W.L., Connors, S., Denton, F., Diongue-Niang, A., Dodman, D., Garschagen, M., Geden, O., Hayward, B., Jones, C., Jotzo, F., Krug, T., Lasco, R., Lee, Y.-Y., Masson-Delmotte, V., Meinshausen, M., Mintenbeck, K., Mokssit, A., Otto, F.E.L., Pathak, M., Pirani, A., Poloczanska, E., Pörtner, H.-O., Revi, A., Roberts, D.C., Roy, J., Ruane, A.C., Skea, J., Shukla, P.R., Slade, R., Slangen, A., Sokona, Y., Sörensson, A.A., Tignor, M., Van Vuuren, D., Wei, Y.-M., Winkler, H., Zhai, P., Zommers, Z., Hourcade, J.-C., Johnson, F.X., Pachauri, S., Simpson, N.P., Singh, C., Thomas, A., Totin, E., Alegría, A., Armour, K., Bednar-Friedl, B., Blok, K., Cissé, G., Dentener, F., Eriksen, S., Fischer, E., Garner, G., Guivarch, C., Haasnoot, M., Hansen, G., Hauser, M., Hawkins, E., Hermans, T., Kopp, R., Leprince-Ringuet, N., Lewis, J., Ley, D., Ludden, C., Niamir, L., Nicholls, Z., Some, S., Szopa, S., Trewin, B., Van Der Wijst, K.-I., Winter, G., Witting, M., Birt, A., Ha, M., 2023. IPCC, 2023: Climate Change 2023: Synthesis Report. Contribution of Working Groups I, II and III to the Sixth Assessment Report of the Intergovernmental Panel on Climate Change [Core Writing Team, H. Lee and J. Romero (eds.)]. IPCC, Geneva, Switzerland. Intergovernmental Panel on Climate Change (IPCC). 10.59327/IPCC/AR6-9789291691647

Campi, P., Palumbo, A.D., Mastrorilli, M., 2009. Effects of tree windbreak on microclimate and wheat productivity in a Mediterranean environment. European Journal of Agronomy 30, 220–227. 10.1016/j.eja.2008.10.004

Chichaghare, A., Chavan, S., Rawale, G., Uthappa, A., Kakade, V., Morade, A., Changan, S., Paul, N., Gawade, V., Khapte, P., Basavaraj, P., Babar, R., Adavi, S.B., Nangare, D., Harisha, C., Halli, H.M., Reddy, K., 2026. Tree shade mitigates stress and enhances chickpea productivity: Insights from an *Emblica officinalis*-based agroforestry system in semi-arid shallow Basaltic Deccan Plateau, India. Trees, Forests and People 24, 101164. 10.1016/j.tfp.2026.101164

Disciglio, G., Stasi, A., Tarantino, A., Frabboni, L., 2025. Microclimate Modification, Evapotranspiration, Growth and Essential Oil Yield of Six Medicinal Plants Cultivated Beneath a Dynamic Agrivoltaic System in Southern Italy. Plants 14, 2428. 10.3390/plants14152428

Dupraz, C., Blitz-Frayret, C., Lecomte, I., Molto, Q., Et., A., 2018. Influence of latitude on the light availability for intercrops in an agroforestry alley-cropping system. Agroforestry Systems. 10.1007/s10457-018-0214-x

El-Zawily, H.M.A., Abo El-Enin, M.M.S., Elmenofy, H.M., Hassan, I.F., Manolikaki, I., Koubouris, G., Alam-Eldein, S.M., 2024. Improving Yield and Quality of ‘Balady’ Mandarin Trees by Using Shading Techniques and Reflective Materials in Response to Climate Change Under Flood Irrigation Conditions. Agronomy 14, 2456. 10.3390/agronomy14112456

Farella, M.M., Fisher, J.B., Jiao, W., Key, K.B., Barnes, M.L., 2022. Thermal remote sensing for plant ecology from leaf to globe. Journal of Ecology 110, 1996–2014. 10.1111/1365-2745.13957

Feldhake, C.M., Belesky, D.P., 2009. Photosynthetically active radiation use efficiency of Dactylis glomerata and Schedonorus phoenix along a hardwood tree-induced light gradient. Agroforest Syst 75, 189–196. 10.1007/s10457-008-9175-9

Fernández, J.E., Cuevas, M.V., 2010. Irrigation scheduling from stem diameter variations: A review. Agricultural and Forest Meteorology. 10.1016/j.agrformet.2009.11.006

Fernández, M.E., Gyenge, J., Licata, J., Schlichter, T., Bond, B.J., 2008. Belowground interactions for water between trees and grasses in a temperate semiarid agroforestry system. Agroforest Syst 74, 185–197. 10.1007/s10457-008-9119-4

Figueiredo, A.C., Barroso, J.G., Pedro, L.G., Scheffer, J.J.C., 2008. Factors affecting secondary metabolite production in plants: volatile components and essential oils. Flavour & Fragrance J 23, 213–226. 10.1002/ffj.1875

Gosme, M., Dufour, L., Aguirre, H.D.I., Dupraz, C., 2016. Microclimatic effect of agroforestry on diurnal temperature cycle 5.

Greis, I., Kellomäki, S., 1981. Crown structure and stem growth of Norway spruce undergrowth under varying shading. Silva Fennica 15.

Herbette, S., Le Menn, A., Rousselle, P., Ameglio, T., Faltin, Z., Branlard, G., Eshdat, Y., Julien, J.-L., Drevet, J.R., Roeckel-Drevet, P., 2005. Modification of photosynthetic regulation in tomato overexpressing glutathione peroxidase. Biochimica et Biophysica Acta (BBA)-General Subjects 1724, 108–118.

Hirasawa, T., Hsiao, T.C., 1999. Some characteristics of reduced leaf photosynthesis at midday in maize growing in the field. Field Crops Research. 10.1016/S0378-4290(99)00005-2

Holtta, T., Makinen, H., Nojd, P., Makela, A., Et., A., 2010. A physiological model of softwood cambial growth. Tree Physiology. 10.1093/treephys/tpq068

Inurreta-Aguirre, H.D., Lauri, P.-É., Dupraz, C., Gosme, M., 2018. Yield components and phenology of durum wheat in a Mediterranean alley-cropping system. Agroforest Syst 92, 961–974. 10.1007/s10457-018-0201-2

Jha, A., Heiser, G., Kelvey, R., Huang, Q., 2026. Crop Yield Responses to Reduced Solar Radiation in Agrivoltaic Systems: Crop-Specific Patterns and Shading Thresholds. Agronomy 16, 985. 10.3390/agronomy16100985

Jing, M., Herbette, S., Vandame, M., Kositsup, B., Kasemsap, P., Cavaloc, E., Julien, J., Améglio, T., Roeckel-Drevet, P., 2009. Effect of chilling on photosynthesis and antioxidant enzymes in Hevea brasiliensis Muell. Arg Trees: Structure and Function 23, 863–874.

Jones, H.G., Rotenberg, E., 2011. Energy, Radiation and Temperature Regulation in Plants, in: Encyclopedia of Life Sciences. John Wiley & Sons, Ltd. 10.1002/9780470015902.a0003199.pub2

Jose, S., 2009. Agroforestry for ecosystem services and environmental benefits: an overview. Agroforest Syst 76, 1–10. 10.1007/s10457-009-9229-7

Juillion, P., Lopez, G., Fumey, D., Lesniak, V., Génard, M., Vercambre, G., 2022. Shading apple trees with an agrivoltaic system: Impact on water relations, leaf morphophysiological characteristics and yield determinants. Scientia Horticulturae 306, 111434. 10.1016/j.scienta.2022.111434

Kabir, M.Y., Nambeesan, S.U., Bautista, J., Díaz-Pérez, J.C., 2022. Plant water status, plant growth, and fruit yield in bell pepper (Capsicum annum L.) under shade nets. Scientia Horticulturae. 10.1016/j.scienta.2022.111241

Kangur, O., Kupper, P., Sellin, A., 2017. Predawn disequilibrium between soil and plant water potentials in light of climate trends predicted for northern Europe. Reg Environ Change 17, 2159–2168. 10.1007/s10113-017-1183-8

Kanzler, M., Böhm, C., Mirck, J., Schmitt, D., Veste, M., 2019. Microclimate effects on evaporation and winter wheat (Triticum aestivum L.) yield within a temperate agroforestry system. Agroforest Syst 93, 1821–1841. 10.1007/s10457-018-0289-4

Karvatte, N., Miyagi, E.S., de Oliveira, C.C., Barreto, C.D., Et., A., 2020. Infrared thermography for microclimate assessment in agroforestry systems. Science of The Total Environment. 10.1016/j.scitotenv.2020.139252

Katsoulis, G.I., Kimbaris, A.C., Anastasaki, E., Damalas, C.A., Kyriazopoulos, A.P., 2022. Chamomile and Anise Cultivation in Olive Agroforestry Systems. Forests 13, 128. 10.3390/f13010128

Kramer, P.J., 1988. Changing concepts regarding plant water relations. Plant, Cell & Environment 11, 565–567. 10.1111/j.1365-3040.1988.tb01796.x

Lalević, D., Ilić, Z.S., Stanojević, L., Milenković, L., Šunić, L., Kovač, R., Kovačević, D., Danilović, B., Milenković, A., Stanojević, J., Cvetković, D., 2023. Shade-Induced Effects on Essential Oil Yield, Chemical Profiling, and Biological Activity in Some Lamiaceae Plants Cultivated in Serbia. Horticulturae 9, 84. 10.3390/horticulturae9010084

Lamacque, L., Charrier, G., Farnese, F.D.S., Lemaire, B., Améglio, T., Herbette, S., 2020. Drought-Induced Mortality: Branch Diameter Variation Reveals a Point of No Recovery in Lavender Species. Plant Physiol. 183, 1638–1649. 10.1104/pp.20.00165

Lammerink, J., Wallace, A.R., Porter, N.G., 1989. Effects of harvest time and postharvest drying on oil from lavandin ( *Lavandula* × *intermedia* ). New Zealand Journal of Crop and Horticultural Science 17, 315–326. 10.1080/01140671.1989.10428051

Lee, J., Kirn, Y., Choi, Y., Ahn, D., 1996. AGRONOMIC FACTORS AFFECTING ROOT YIELD AND ESSENTIAL OIL CONTENTS OF VALERIANA FAURIEI VAR. DASYCARPA HARA AND V. OFFICINALIS L. IN KOREA. Acta Horticulturae. 10.17660/ActaHortic.1996.426.57

Leigh, A., Sevanto, S., Close, J. d., Nicotra, A. b., 2017. The influence of leaf size and shape on leaf thermal dynamics: does theory hold up under natural conditions? Plant, Cell & Environment 40, 237–248. 10.1111/pce.12857

Lis-Balchin, M. (Ed.), 2002. Lavender: the genus lavandula, 1. publ. ed, Medicinal and aromatic plants - industrial profiles. Taylor & Francis, London.

Livesley, S.J., Gregory, P.J., Buresh, R.J., 2004. Competition in tree row agroforestry systems. 3. Soil water distribution and dynamics. Plant and Soil 264, 129–139. 10.1023/B:PLSO.0000047750.80654.d5

Lopes, M.J. dos S., Dias-Filho, M.B., Neto, M.A.M., Cruz, E.D., 2015. Morphophysiological Behavior and Cambial Activity in Seedlings of Two Amazonian Tree Species under Shade. Journal of Botany 2015, 863968. 10.1155/2015/863968

López-Díaz, M.L., Benítez, R., Rolo, V., Moreno, G., 2026. How cereal acclimatation under tree canopy affects grain production in Mediterranean agroforestry systems. Agroforest Syst 100, 115. 10.1007/s10457-026-01435-5

Lott, J.E., Ong, C.K., Black, C.R., 2009. Understorey microclimate and crop performance in a Grevillea robusta-based agroforestry system in semi-arid Kenya. Agricultural and Forest Meteorology 149, 1140–1151. 10.1016/j.agrformet.2009.02.002

Lu, C., Zhang, J., 1999. Effects of water stress on photosystem II photochemistry and its thermostability in wheat plants. J Exp Bot 50, 1199–1206. 10.1093/jxb/50.336.1199

Lubbe, A., Verpoorte, R., 2011. Cultivation of medicinal and aromatic plants for specialty industrial materials. Industrial Crops and Products 34, 785–801. 10.1016/j.indcrop.2011.01.019

Magarelli, A., Mazzeo, A., Alhajj Ali, S., Ferrara, G., 2025. Shading enhanced microclimate variability, photomorphogenesis and yield components in a grapevine agrivoltaic system in semi-arid Mediterranean conditions in Puglia region, southeastern Italy. Scientia Horticulturae 350, 114311. 10.1016/j.scienta.2025.114311

Magarelli, A., Mazzeo, A., Ferrara, G., 2024. Fruit Crop Species with Agrivoltaic Systems: A Critical Review. Agronomy 14, 722. 10.3390/agronomy14040722

Mahmood, A., Hu, Y., Tanny, J., Asante, E.A., 2018. Effects of shading and insect-proof screens on crop microclimate and production: A review of recent advances. Scientia Horticulturae 241, 241–251. 10.1016/j.scienta.2018.06.078

Mambrí, A.P., Andriolo, J.L., Manfron, M.P., Pinheiro, S.M., Cardoso, F.L., Neves, M.G., 2018. Yield and composition of lavender essential oil grown in substrate. Hortic. Bras. 36, 259–264. 10.1590/s0102-053620180219

Mantino, A., Tozzini, C., Bonari, E., Mele, M., Ragaglini, G., 2021. Competition for Light Affects Alfalfa Biomass Production More Than Its Nutritive Value in an Olive-Based Alley-Cropping System. Forests 12, 233. 10.3390/f12020233

Marcelis, L.F.M., 1996. Sink strength as a determinant of dry matter partitioning in the whole plant. Journal of Experimental Botany. 10.1093/jxb/47.Special_Issue.1281

Marrou, H., Guilioni, L., Dufour, L., Dupraz, C., Wery, J., 2013. Microclimate under agrivoltaic systems: Is crop growth rate affected in the partial shade of solar panels? Agricultural and Forest Meteorology 177, 117–132. 10.1016/j.agrformet.2013.04.012

Marthe, F., 2020. Petroselinum crispum (Mill.) Nyman (Parsley), in: Novak, J., Blüthner, W.-D. (Eds.), Medicinal, Aromatic and Stimulant Plants, Handbook of Plant Breeding. Springer International Publishing, Cham, pp. 435–466. 10.1007/978-3-030-38792-1_13

Maxwell, K., Johnson, G.N., 2000. Chlorophyll fluorescence—a practical guide. Journal of Experimental Botany. 10.1093/jexbot/51.345.659

Mira-García, A.B., Conejero, W., Vera, J., Ruiz-Sánchez, M.C., 2022. Water status and thermal response of lime trees to irrigation and shade screen. Agricultural Water Management 272, 107843. 10.1016/j.agwat.2022.107843

Montanaro, G., Dichio, B., Xiloyannis, C., 2009. Shade mitigates photoinhibition and enhances water use efficiency in kiwifruit under drought. Photosynthetica 47, 363–371. 10.1007/s11099-009-0057-9

Monteith, J.L., Ong, C.K., Corlett, J.E., 1991. Microclimatic interactions in agroforestry systems. Forest Ecology and Management 45, 31–44. 10.1016/0378-1127(91)90204-9

Monteith, J.L., Unsworth, M.H., 2008. Principles of environmental physics, 3rd ed. ed. Elsevier, Amsterdam ; Boston.

Moradkhani, H., Sargsyan, E., Bibak, H., Naseri, B., Sadat-Hosseini, M., Fayazi-Barjin, A., Meftahizade, H., 2010. Melissa officinalis L., a valuable medicine plant: A review. Journal of Medicinal Plants Research 4, 2753–2759.

Mupambi, G., Anthony, B.M., Layne, D.R., Musacchi, S., Serra, S., Schmidt, T., Kalcsits, L.A., 2018. The influence of protective netting on tree physiology and fruit quality of apple: A review. Scientia Horticulturae 236, 60–72. 10.1016/j.scienta.2018.03.014

Murray, F.W., 1967. On the Computation of Saturation Vapor Pressure. Journal of Applied Meteorology and Climatology 6, 203–204. 10.1175/1520-0450(1967)006%3C0203:OTCOSV%3E2.0.CO;2

Muschler, R.G., 2016. Agroforestry: Essential for Sustainable and Climate-Smart Land Use?, in: Pancel, L., Köhl, M. (Eds.), Tropical Forestry Handbook. Springer Berlin Heidelberg, Berlin, Heidelberg, pp. 2013–2116.

Narjesi, V., Moghadam, J.F., Ghasemi-Soloklui, A.A., 2023. Effects of Photo-selective Shade Net Color and Shading Percentage on Reducing Sunburn and Increasing the Quantity and Quality of Pomegranate Fruit.

Nogues, S., Munne-Bosch, S., Casadesus, J., Lopez-Carbonell, M., Alegre, L., 2001. Daily time course of whole-shoot gas exchange rates in two drought-exposed Mediterranean shrubs. Tree Physiology 21, 51–58. 10.1093/treephys/21.1.51

Ortuño, M.F., Conejero, W., Moreno, F., Moriana, A., Intrigliolo, D.S., Biel, C., Mellisho, C.D., Pérez-Pastor, A., Domingo, R., Ruiz-Sánchez, M.C., Casadesus, J., Bonany, J., Torrecillas, A., 2010. Could trunk diameter sensors be used in woody crops for irrigation scheduling? A review of current knowledge and future perspectives. Agricultural Water Management 97, 1–11. 10.1016/j.agwat.2009.09.008

Palada, M., Becker, B.N., Mitchell, J.M., Nair, P.K.R., 2004. Cultivation of medicinal plants in alley cropping system with Moringa oleifera in the Virgin Islands, in: Annual Agriculture and Food Fair of the U.S. Virgin Islands.

Pallotti, L., Silvestroni, O., Dottori, E., Lattanzi, T., Lanari, V., 2023. Effects of shading nets as a form of adaptation to climate change on grapes production: a review. 10.20870/oeno-one.2023.57.2.7414

Pandey, A.K., Kumar, P., Saxena, M.J., Maurya, P., 2020. Distribution of aromatic plants in the world and their properties. Feed Additives. 10.1016/B978-0-12-814700-9.00006-6

Pang, K., Van Sambeek, J.W., Navarrete-Tindall, N.E., Lin, C.-H., Jose, S., Garrett, H.E., 2019. Responses of legumes and grasses to non-, moderate, and dense shade in Missouri, USA. I. Forage yield and its species-level plasticity. Agroforest Syst 93, 11–24. 10.1007/s10457-017-0067-8

Pardhi, D.H., Mane, Dr.V., Bisane, R.D., 2020. Incident photosyntehticaly active radiation under and outside the canopy of Acacia nilotica (Linn.) for agroforestry system. Int. J. Chem. Stud. 8, 1973–1976. 10.22271/chemi.2020.v8.i6ab.11056

Patočka, J., Jakl, J., 2010. Biomedically relevant chemical constituents of Valeriana officinalis. J Appl Biomed 8, 11–18. 10.2478/v10136-009-0002-z

Peng, R., Xu, H., Bi, H., Wang, N., 2025. Accumulated Photosynthetically Active Radiation and Its Heterogeneity Collectively Decrease Soybean Yield in Apple-Based Intercropping Systems. Agronomy 15, 581. 10.3390/agronomy15030581

Pokajewicz, K., Czarniecka-Wiera, M., Krajewska, A., Maciejczyk, E., Wieczorek, P.P., 2023. Lavandula × intermedia—A Bastard Lavender or a Plant of Many Values? Part I. Biology and Chemical Composition of Lavandin. Molecules 28, 2943. 10.3390/molecules28072943

Poorter, H., Niinemets, Ü., Poorter, L., Wright, I.J., Villar, R., 2009. Causes and consequences of variation in leaf mass per area (LMA): a meta analysis. New Phytologist 182, 565–588. 10.1111/j.1469-8137.2009.02830.x

Poorter, H., Niklas, K.J., Reich, P.B., Oleksyn, J., Poot, P., Mommer, L., 2012. Biomass allocation to leaves, stems and roots: meta-analyses of interspecific variation and environmental control. New Phytologist 193, 30–50. 10.1111/j.1469-8137.2011.03952.x

Querné, A., Battie-laclau, P., Dufour, L., Wery, J., Dupraz, C., 2017. Effects of walnut trees on biological nitrogen fixation and yield of intercropped alfalfa in a Mediterranean agroforestry system. European Journal of Agronomy 84, 35–46. 10.1016/j.eja.2016.12.001

Quinkenstein, A., Wöllecke, J., Böhm, C., Grünewald, H., Freese, D., Schneider, B.U., Hüttl, R.F., 2009. Ecological benefits of the alley cropping agroforestry system in sensitive regions of Europe. Environmental Science & Policy, Sustainability impact assessment and land-use policies for sensitive regions 12, 1112–1121. 10.1016/j.envsci.2009.08.008

Rao, M.R., Palada, M.C., Becker, B.N., 2004. Medicinal and aromatic plants in agroforestry systems. Agroforestry Systems 61, 107–122. 10.1023/B:AGFO.0000028993.83007.4b

Rezaei, E.E., Webber, H., Asseng, S., Boote, K., Durand, J.L., Ewert, F., Martre, P., MacCarthy, D.S., 2023. Climate change impacts on crop yields. Nat Rev Earth Environ 4, 831–846. 10.1038/s43017-023-00491-0

Sabry, R.M., Kandil, M.A.M., Ahmed, S.S., 2016. Growth and Quality of Sage (Salvia officinalis), Parsley (Petroselinum crispum) and Nasturtium (Tropaeolum majus) as Affected by Water Deficit.

Saunier, A., Ormeño, E., Moja, S., Fernandez, C., Robert, E., Dupouyet, S., Despinasse, Y., Baudino, S., Nicolè, F., Bousquet-Mélou, A., 2022. Lavender sensitivity to water stress: Comparison between eleven varieties across two phenological stages. Industrial Crops and Products 177, 114531. 10.1016/j.indcrop.2022.114531

Schindelin, J., Arganda-Carreras, I., Frise, E., Kaynig, V., Longair, M., Pietzsch, T., Preibisch, S., Rueden, C., Saalfeld, S., Schmid, B., Tinevez, J.-Y., White, D.J., Hartenstein, V., Eliceiri, K., Tomancak, P., Cardona, A., 2012. Fiji: an open-source platform for biological-image analysis. Nat Methods 9, 676–682. 10.1038/nmeth.2019

Scholander, P.F., Bradstreet, E.D., Hemmingsen, E.A., Hammel, H.T., 1965. Sap Pressure in Vascular Plants. Science 148, 339–346. 10.1126/science.148.3668.339

Sida, T.S., Baudron, F., Kim, H., Giller, K.E., 2018. Climate-smart agroforestry: *Faidherbia albida* trees buffer wheat against climatic extremes in the Central Rift Valley of Ethiopia. Agricultural and Forest Meteorology 248, 339–347. 10.1016/j.agrformet.2017.10.013

Singh, K., Rajeswara Rao, B.R., Singh, C.P., Bhattacharya, A.K., Kaul, P.N., 1998. Production potential of aromatic crops in the alleys of Eucalyptus citriodora in semi - arid tropical climate of south India. Journal of Medicinal and Aromatic Plant Sciences 20, 749–752.

Temani, F., Bouaziz, A., Daoui, K., Wery, J., Barkaoui, K., 2021. Olive agroforestry can improve land productivity even under low water availability in the South Mediterranean. Agriculture, Ecosystems & Environment 307, 107234. 10.1016/j.agee.2020.107234

Tezcan, N.Y., Taşpinar, H., Korkmaz, C., 2022. Effects of Shade Nets on The Microclimate and Growth of Tomato. J Agr Sci-Tarim Bili. 10.15832/ankutbd.1073156

Touil, S., Richa, A., Fizir, M., Bingwa, B., 2021. Shading effect of photovoltaic panels on horticulture crops production: a mini review. Reviews in Environmental Science and Bio/Technology. 10.1007/s11157-021-09572-2

Ukwu, U.N., Muller, O., Meier-Grüll, M., Uguru, M.I., 2025. Agrivoltaics shading enhanced the microclimate, photosynthesis, growth and yields of vigna radiata genotypes in tropical Nigeria. Sci Rep 15, 1190. 10.1038/s41598-024-84216-3

Valladares, F., Niinemets, Ü., 2008. Shade Tolerance, a Key Plant Feature of Complex Nature and Consequences. Annu. Rev. Ecol. Evol. Syst. 39, 237–257. 10.1146/annurev.ecolsys.39.110707.173506

Van Baalen, J., Ernst, W.H.O., Van Andel, J., Janssen, D.W., Nelissen, H.J.M., 1990. Reproductive allocation in plants of *Scrophularia nodosa* grown at various levels of irradiance and soil fertility. Acta Botanica Neerlandica 39, 183–196. 10.1111/j.1438-8677.1990.tb01486.x

Vogel, S., 2009. Leaves in the lowest and highest winds: temperature, force and shape. New Phytologist 183, 13–26. 10.1111/j.1469-8137.2009.02854.x

Von Arx, G., Dobbertin, M., Rebetez, M., 2012. Spatio-temporal effects of forest canopy on understory microclimate in a long-term experiment in Switzerland. Agricultural and Forest Meteorology 166-167, 144–155. 10.1016/j.agrformet.2012.07.018

Wang, J., Xu, Z., 2026. Plant response to and recovery from drought. Current Biology 36, R343–R362. 10.1016/j.cub.2026.02.025

Weselek, A., Bauerle, A., Hartung, J., Zikeli, S., Lewandowski, I., Högy, P., 2021. Agrivoltaic system impacts on microclimate and yield of different crops within an organic crop rotation in a temperate climate. Agron. Sustain. Dev. 41, 59. 10.1007/s13593-021-00714-y

Xiao, J., Fisher, J.B., Hashimoto, H., Ichii, K., Parazoo, N.C., 2021. Emerging satellite observations for diurnal cycling of ecosystem processes. Nat. Plants 7, 877–887. 10.1038/s41477-021-00952-8

Yasuda, Y., Utsumi, Y., Tan, X., Tashiro, N., Fukuda, K., Koga, S., 2018. Suppression of growth and death of meristematic tissues in Abies sachalinensis under strong shading: comparisons between the terminal bud, the terminally lateral bud and the stem cambium. J Plant Res 131, 817–825. 10.1007/s10265-018-1051-8

Zha, Q., Yin, X., Xi, X., Jiang, A., 2022. Colored Shade Nets Can Relieve Abnormal Fruit Softening and Premature Leaf Senescence of “Jumeigui” Grapes during Ripening under Greenhouse Conditions. Plants 11, 1227. 10.3390/plants11091227

Zhao, W., Qualls, R.J., Berliner, P.R., 2003. Modeling of the short wave radiation distribution in an agroforestry system. Agricultural and Forest Meteorology 118, 185–206. 10.1016/S0168-1923(03)00108-4

Ziegler, Y., Grote, R., Alongi, F., Knüver, T., Ruehr, N.K., 2024. Capturing drought stress signals: the potential of dendrometers for monitoring tree water status. Tree Physiology 44, tpae140. 10.1093/treephys/tpae140

Zubay, P., Ruttner, K., Ladányi, M., Deli, J., Németh Zámboriné, É., Szabó, K., 2021. In the shade – Screening of medicinal and aromatic plants for temperate zone agroforestry cultivation. Industrial Crops and Products 170, 113764. 10.1016/j.indcrop.2021.113764

